# Genome-wide functional screen of 3’UTR variants uncovers causal variants for human disease and evolution

**DOI:** 10.1101/2021.01.13.424697

**Authors:** Dustin Griesemer, James R Xue, Steven K Reilly, Jacob C Ulirsch, Kalki Kukreja, Joe Davis, Masahiro Kanai, David K Yang, Stephen B Montgomery, Carl D Novina, Ryan Tewhey, Pardis C Sabeti

## Abstract

3’ untranslated region (3’UTR) variants are strongly associated with human traits and diseases, yet few have been causally identified. We developed the Massively Parallel Reporter Assay for 3’UTRs (MPRAu) to sensitively assay 12,173 3’UTR variants. We applied MPRAu to six human cell lines, focusing on genetic variants associated with genome-wide association studies (GWAS) and human evolutionary adaptation. MPRAu expands our understanding of 3’UTR function, suggesting that low-complexity sequences predominately explain 3’UTR regulatory activity. We adapt MPRAu to uncover diverse molecular mechanisms at base-pair resolution, including an AU-rich element of *LEPR* linked to potential metabolic evolutionary adaptations in East Asians. We nominate hundreds of 3’UTR causal variants with genetically fine-mapped phenotype associations. Using endogenous allelic replacements, we characterize one variant that disrupts a miRNA site regulating the viral defense gene *TRIM14*, and one that alters *PILRB* abundance, nominating a causal variant underlying transcriptional changes in age-related macular degeneration.

## Introduction

Over the past two decades, thousands of variant-trait associations have been identified by genome-wide association studies (GWAS) (Buniello et al., 2019). However, GWAS have been hindered in elucidating the mechanisms of complex disease by two limitations: (1) linkage disequilibrium (LD) causes neighboring neutral polymorphisms to display similarly strong associations with causal alleles, greatly increasing the experimental burden for functional validation; (2) over 90% of associations reside in non-coding regions of the genome (Gusev et al., 2014; Maurano et al., 2012), where functional interpretation is much more difficult than in coding regions.

3’ untranslated regions (3’UTRs) contain a particularly important class of noncoding variants that can impact post-transcriptional and translational processes. Causal peripheral blood cis-expression Quantitative Trait Loci (eQTL) variants are 4-fold enriched to be in 3’UTRs, a level matching that of promoter elements (Wang et al., 2020b). Across all tissues in the Genotype-Tissue Expression project (GTEx), eQTLs in 3’UTRs are found to be 2-fold enriched, the largest enrichment amongst all non-coding regions (The GTEx Consortium, 2020). Untranslated regions harbor the largest enrichment of GWAS heritability (5-fold) of all non-coding categories except for transcription start sites, emphasizing a substantial role for post-transcriptional activities in human regulatory variation (Finucane et al., 2015).

Although 3’UTR variants are crucial to understanding human phenotypic variation, only a handful of causal 3’UTR variants have been described. They include BAFF-var in *TNFSF13B* associated with lupus and multiple sclerosis (Steri et al., 2017), rs13702 in *LPL* associated with HDL-cholesterol levels (Richardson et al., 2013), and rs12190287 in *TCF21* associated with coronary artery disease (Miller et al., 2014). Each case required meta-analysis across multiple trait or population datasets and annotation with well-known regulatory factors, before being pursued with low-throughput luciferase confirmations. These factors demonstrate how current 3’UTR causal variant discovery is burdensome and highlight the need for high-throughput tools to characterize the functional impact of 3’UTR variants on gene expression.

The development of Massively Parallel Reporter Assays (MPRAs) has enabled simultaneous testing of thousands of variants for cis-regulatory activity to nominate causal variants in non-coding regions (van Arensbergen et al., 2019; Choi et al., 2020; Kircher et al., 2019; Klein et al., 2019; Liu et al., 2017; Sample et al., 2019; Tewhey et al., 2016; Ulirsch et al., 2016), Historically MPRA has been primarily applied to understand transcriptional regulation. Several studies have adapted MPRA to test 3’UTR sequences but to date, none have comprehensively characterized the impact of genetic variation in 3’UTRs (Bogard et al., 2019; Litterman et al., 2019; Oikonomou et al., 2014; Siegel et al., 2020; Vainberg Slutskin et al., 2018, 2019; Zhao et al., 2014).

Here, we developed the Massively Parallel Reporter Assay for 3’UTRs (MPRAu) to quantify allelic expression differences for thousands of 3’UTR variants simultaneously in a high-throughput, accurate, and reproducible manner. MPRAu detects distinct aspects of 3’UTR regulation, allowing us to understand general sequence features governing transcript abundance via computational modeling, pinpoint exact sequence architectures underlying variant functionality including RNA structure and RNA-binding protein (RBP) occupancy, and nominate causal variants. We utilize MPRAu to comprehensively test disease-associated, as well as evolutionarily adaptive, 3’UTR genetic variation in six human cell lines. From our functionally nominated causal variants, we also more deeply characterize two variants using CRISPR-induced allelic replacement.

## Results

### MPRAu reproducibly characterizes the functions of thousands of 3’UTR elements

We applied MPRAu to systematically evaluate the functional effects of genetic variation from 3’UTRs. To do so, we designed and synthesized 100 base pair (bp) oligonucleotides derived from human 3’UTRs, centered on, and differing only with respect to the variant’s ‘reference’ (ref) or ‘alternate’ (alt) alleles (**Fig. 1a**), for testing using MPRAu. We cloned the oligo pool into the 3’UTR of a plasmid reporter gene controlled by a moderately strong promoter. By transfecting our pool into cell lines of interest, and sequencing both the plasmid pool and mRNA from cells, we could compare steady-state RNA expression effects of each 3’UTR oligonucleotide (from either differential mRNA decay or transcription). We refer to 3’UTR oligo backgrounds that increase mRNA levels as having ‘augmenting’ effects and those that decrease transcript levels as having ‘attenuating’ effects. We also quantify differences between sequences bearing the ref versus alt allele and refer to alleles with a statistically significant ‘allelic skew’ as transcript abundance-modulating variants (tamVars). In addition, MPRAu employs several quality controls to minimize bias, including using random barcodes to ensure adequate library complexity (Methods).

**Figure 1:**
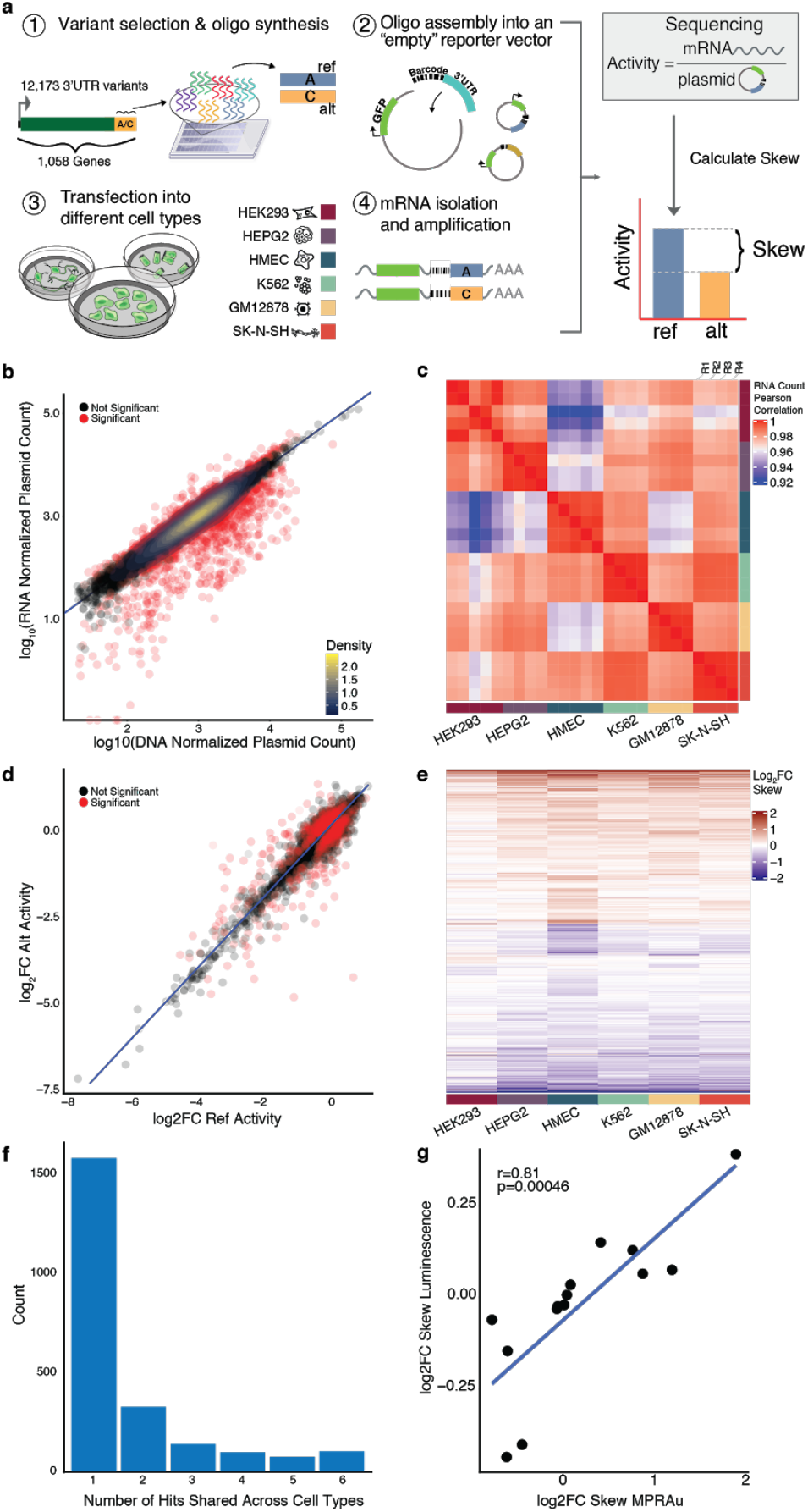
MPRAu actively and reproducibly recapitulates known 3’UTR activity. **a**, Overview of MPRAu: (1) Synthesis of oligonucleotides across variants of interest. (2) Oligos are PCR-amplified and inserted into a vector backbone downstream of GFP and adjacent to a random hexamer barcode. (3) The vector pool is transfected into cell lines of interest, (4) mRNA extraction and sequencing. mRNA sequencing counts (4) are compared to plasmid counts (2) to determine the relative expression of ref and alt alleles. **b**, Scatterplot of HEK293 RNA normalized counts plotted against DNA normalized counts demonstrates that most oligos with significant activity show attenuating effects. **c**, Heatmap of the pairwise correlation of RNA counts across all replicates demonstrates strong replicate concordance. **d**, tamVars (red) are variants that have a significantly different alt activity compared with ref activity (data for HEK293 plotted). **e**, log2FC allelic skews for all variants with significant effects in at least one cell type across all tested cell types. Each row specifies a tamVar. **f**, Barplot depicting tamVar sharing across one to all six cell types. **g**, tamVar allelic skews are reproducible via low-throughput luciferase assays.

We applied MPRAu to identify functional 3’UTR variants associated with human disease and evolutionary selection, testing 12,173 3’UTR variants. As the causal variant (s) underlying human traits and diseases can be amongst many variants associated with GWAS tag SNPs, we tested 3’UTR SNPs and insertion/deletions (indels) (minor allele frequency (MAF)≧5%) in LD with tag SNPs (minimum r^2^=0.8) from the NHGRI-EBI GWAS catalog (Welter et al., 2014), totaling 2,153 putative disease-associated variants from 1,556 independent association loci (**Supplementary Table 1**). We also incorporated a set of 9,325 3’UTR SNPs and indels overlapping regions identified as being under positive selection in humans (Grossman et al., 2013) (**Supplementary Table 1**). We also included a set of 46 rare 3’UTR variants (minor allele frequency (MAF) ≤0.01 in Europeans) that are in genes with outlier expression signatures across tissues in the Genotype-Tissue Expression (GTEx) project, which are known to have potential deleterious consequences (Li et al., 2017) (**Supplementary Table 1**). Lastly, across all tested variants, 3,055 were also tested under alternative allelic backgrounds to account for the potential effect of surrounding sequence variants. As genetic variants impacting traits can have tissue-specific effects (GTEx Consortium, 2017; Marbach et al., 2016; Parker et al., 2013), we characterized these variants across six diverse human cellular lines: HEK293 (embryonic kidney), HepG2 (hepatocellular carcinoma), GM12878 (lymphoblastoid), SK-N-SH (neuroblastoma), K562 (leukemia), and a primary cell line (HMEC, mammary epithelial).

**Table 1:**
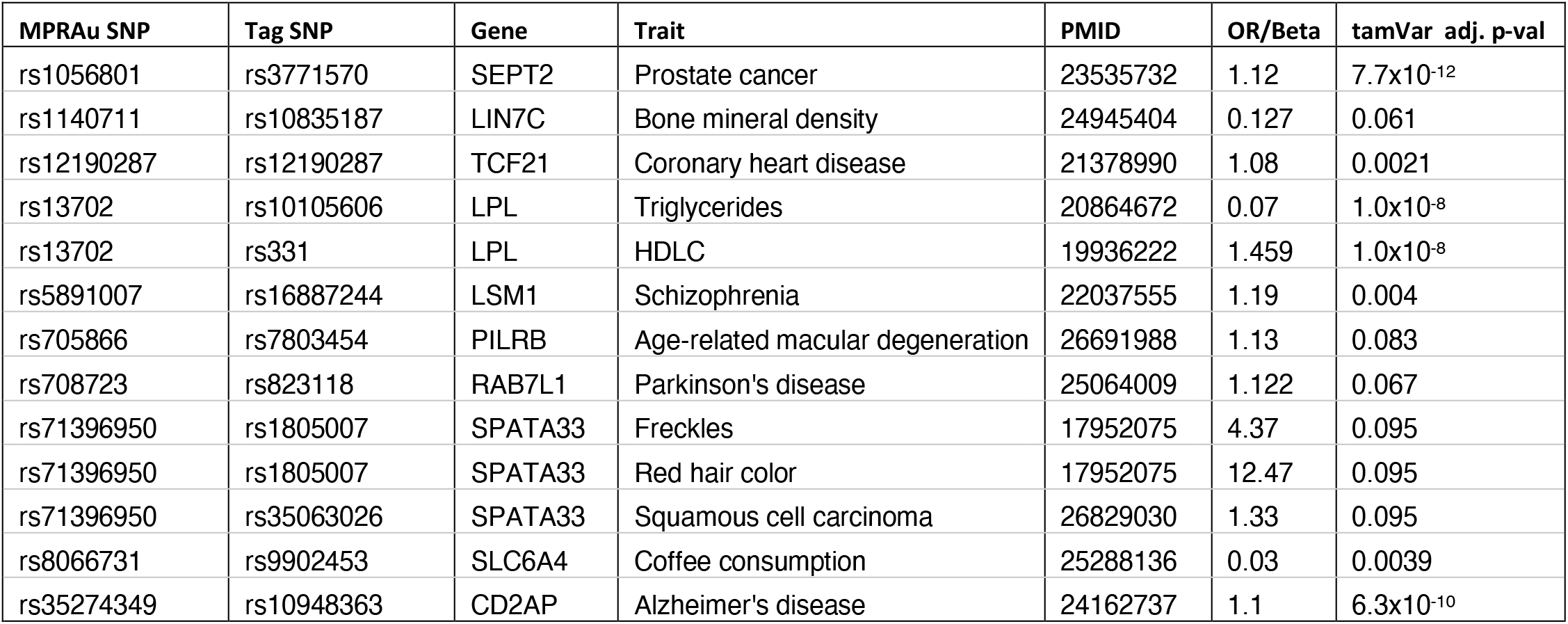
GWAS variants nominated for functional causality by MPRAu. (Most significant adjusted p-value from MPRAu screen is shown)

| MPRAu SNP | Tag SNP | Gene | Trait | PMID | OR/Beta | tamVar adj. p-val |
| --- | --- | --- | --- | --- | --- | --- |
| rs1056801 | rs3771570 | SEPT2 | Prostate cancer | 23535732 | 1.12 | $7.7 \times 10^{-12}$ |
| rs1140711 | rs10835187 | LIN7C | Bone mineral density | 24945404 | 0.127 | 0.061 |
| rs12190287 | rs12190287 | TCF21 | Coronary heart disease | 21378990 | 1.08 | 0.0021 |
| rs13702 | rs10105606 | LPL | Triglycerides | 20864672 | 0.07 | $1.0 \times 10^{-8}$ |
| rs13702 | rs331 | LPL | HDLC | 19936222 | 1.459 | $1.0 \times 10^{-8}$ |
| rs5891007 | rs16887244 | LSM1 | Schizophrenia | 22037555 | 1.19 | 0.004 |
| rs705866 | rs7803454 | PILRB | Age-related macular degeneration | 26691988 | 1.13 | 0.083 |
| rs708723 | rs823118 | RAB7L1 | Parkinson's disease | 25064009 | 1.122 | 0.067 |
| rs71396950 | rs1805007 | SPATA33 | Freckles | 17952075 | 4.37 | 0.095 |
| rs71396950 | rs1805007 | SPATA33 | Red hair color | 17952075 | 12.47 | 0.095 |
| rs71396950 | rs35063026 | SPATA33 | Squamous cell carcinoma | 26829030 | 1.33 | 0.095 |
| rs8066731 | rs9902453 | SLC6A4 | Coffee consumption | 25288136 | 0.03 | 0.0039 |
| rs35274349 | rs10948363 | CD2AP | Alzheimer's disease | 24162737 | 1.1 | $6.3 \times 10^{-10}$ |

We first sought to ensure that our assay was reproducibly capturing expected 3’UTR biological effects. Consistent with the dominant regulatory function of 3’UTRs to attenuate transcript expression, the predominant effect across all MPRAu-tested 3’UTRs was to decrease mRNA abundance (**Fig. 1b**). This effect is reproducibly observed in the strong correlation of normalized RNA read counts between experimental replicates across all cell types (average Pearson correlation (corr)=0.99) (**Fig. 1c**).

Confident in our assay’s ability to assess oligos with regulatory activity, we then identified tamVars altering 3’UTR functionality by comparing expression changes between alleles of the same 3’UTR (using as a threshold a Benjamini-Hochberg adjusted p-value (BH p-adj)<0.1) (**Fig. 1d,e**). We found 2,368 tamVars in total across all cell types, with HMEC having the largest number of tamVars (1,430), and K562 having the least (471), potentially driven by power differences in tamVar detection across different cell types (**Supplementary Table 1**). 104 tamVars were shared across every tested cell line (**Fig. 1f**). Out of the 3,055 variants tested with alternative allelic backgrounds, only 10 tamVars displayed function dependent on its allelic background. To confirm that 3’UTR tamVar effects were reflected in protein levels, we utilized orthogonal methods to measure allelic effects. We performed polysome profiling on HEK293 cells transfected with a subset of our tested library and found polysomal RNA expression highly correlated with steady-state RNA expression (Pearson corr=0.94, p=2.68×10^−11,546^) (**Supplementary Fig. 1a**). In addition, variant effects between polysomal and steady-state RNA were concordant (Pearson corr=0.97, p=1.4×10^−106^) (**Supplementary Fig. 1b**), recapitulating previous findings that steady-state RNA levels are a good proxy for protein levels (Oikonomou et al., 2014; Zhao et al., 2014). To demonstrate that variant effects are directly translatable to the protein level, we also tested a subset of tamVars and non-tamVar controls via luciferase assays (**Supplementary Table 2**). We observed a strong correlation between luciferase luminescence assays and the MPRAu-measure tamVar skew (Pearson corr=0.81, p=4.6×10^−4^) (**Fig. 1g**), which also highlights MPRAu’s ability to triage 3’UTR variant effects over low-throughput luciferase approaches.

### MPRAu sensitively detects 3’UTR regulators and functional sequence variants

To confirm that MPRAu effects are consistent with molecular mechanisms underlying 3’UTR biology, we analyzed features across the entire oligo sequence. We found that GC content and secondary structure, as measured by the predicted minimal free energy, positively correlated with the level of attenuation (percent GC: Pearson corr=-0.24, p=6.59×10^−384^, mfe: Pearson corr=0.27, p=8.87×10^−485^) (**Fig. 2a**). This finding may be explained by the role of high GC content and structuredness in RBP occupancy and therefore functionality (Dominguez et al., 2018; Litterman et al., 2019).

**Figure 2:**
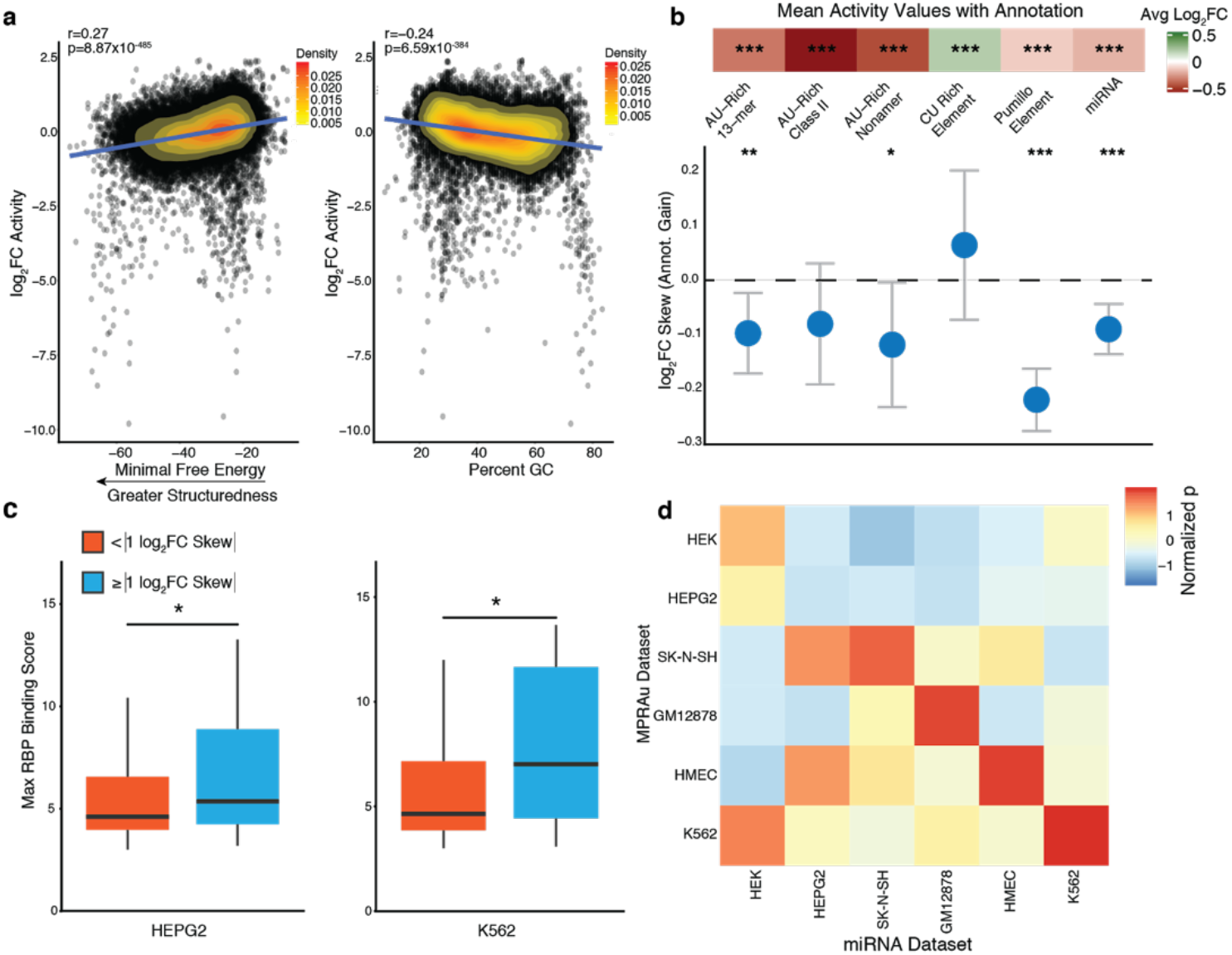
Functional 3’UTR elements overlap known 3’UTR annotations. **a**, Scatterplot showing oligos that are highly structured (more negative minimum free energy, left) and oligos with higher percent GC (right) have greater attenuating effects. HMEC log2FC activity data is representative and plotted. **b, c** (*** p-value<0.001, ** p-value<0.01, * p-value<0.05, tests were performed via a two-sided Wilcoxon rank-sum test). **b**, (top panel) 3’UTR activity shows the expected functional effect directionality from tested sequences that overlap known 3’UTR attenuating annotations (AU-rich, Pumillo, and miRNA) and a known augmenting annotation (CU-rich Element). (bottom panel) Barplot of the allelic skews for variants that acquire 3’UTR annotations of the class listed above demonstrates the expected directional activity gain. Data from all tested cell types were aggregated to produce the plot. **c**, Variants with high allelic skew (| log2FC Skew |≥1) have greater eCLIP RBP binding scores for both HepG2s and K562s than variants with a lesser allelic skew, highlighting the ability of MPRAu to find variants impacting in vivo signatures. **d**, Across 4 cell types (K562, HMEC, GM12878, SK-N-SH), 3’UTR activity was most significantly attenuated (*t*-test) when subsetting on elements with binding sites of the most expressed cell-type-matched miRNAs.

At the finer sequence level, MPRAu captured the expected attenuation and augmentation effects from the presence of empirical RBP motifs (Bakheet et al., 2001; Holcik and Liebhaber, 1997; White et al., 2001) and predicted miRNA motifs (Friedländer et al., 2012; Friedman et al., 2009). Regulatory signatures such as AU-rich elements, the canonical Pumilio motif, and miRNAs motifs exhibited expected attenuating effects on expression (avg log_2_ fold change (log_2_FC) range −0.55 to −0.12, two-sided *t*-test p<1×10^−30^ for all attenuating factors) (**Fig. 2b**). Conversely, CU-rich elements demonstrated their expected augmenting effects on expression (avg log_2_FC=0.18, two-sided *t*-test p=1.58×10^−14^) (**Fig. 2b**). Variants perturbing these predicted elements abrogate the functional effect (two-sided *t*-test p<0.05 for all factors except for CU Rich Element and AU-Rich Class II, but skew directionality is consistent for the latter two) (**Fig. 2b**). Highlighting MPRAu’s ability to capture endogenous perturbation effects, variants with high allelic skews (| log_2_FC Skew |≥1) also are predominantly in regions with elevated in vivo RBP occupancy signatures (Van Nostrand et al., 2016) (two-sided Wilcoxon rank-sum test p=0.02, 0.011, for HepG2 and K562, respectively) (**Fig. 2c**). Furthermore, attenuation of transcript levels in a cell line is generally well explained by its top 10 expressed miRNAs, demonstrating the capacity of MPRAu to capture cell-type-specific effects (**Fig. 2d**). Lastly, an orthogonal analysis of the barcodes in our MPRAu design also validated the effects of known 3’UTR RBP motifs and provides a valuable resource of potential functional hexamers (Methods, **Supplementary Fig. 2, 3, Supplementary Table 3**). Together these data demonstrate the ability of MPRAu to detect the effects of known 3’UTR regulators and cell-type-specific regulatory mechanisms.

### Computational modeling uncovers known and novel features of 3’UTR regulation

Having identified key 3’UTR features underlying MPRAu transcript levels, we trained predictive models for our tested sequences and compared the sensitivity and specificity of several classification models to predict 3’UTR elements with attenuated activity (< −0.5 3’UTR log_2_FC) (Methods). We used 34 total simple, sequence-specific annotations in our initial models, including features derived from nucleotide and dinucleotide composition, homopolymer content, and secondary structure (Methods). Our best model performed well in all cell types, with an average precision of 0.23-0.48 and an area under the receiver operating characteristic curve of 0.67-0.79 (**Fig. 3a,b, Supplementary Fig. 4a-h**). We also used the same features to predict augmenting expression (>0.5 log_2_FC) and found comparable performance (**Supplementary Fig. 5a,b**).

**Figure 3:**
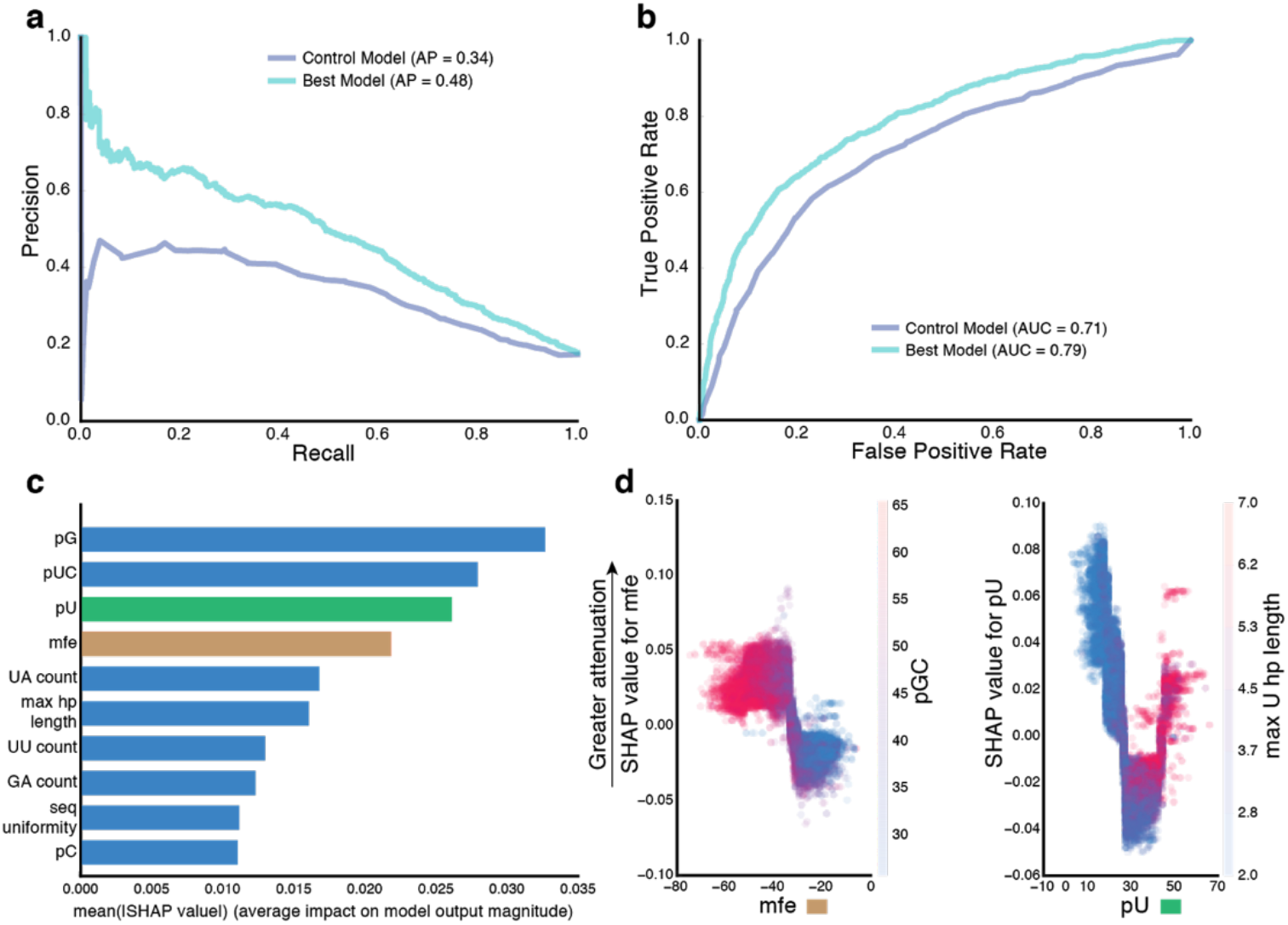
Computational modeling of 3’UTR activity builds accurate predictive models and uncovers features important for prediction. Precision-recall (**a**) and receiver operating characteristic curve (**b**) for the best parameters trained on predicting 3’UTR activity attenuation using the best model (xgboost) (data displayed is for HMEC). The results from a control model (one-variable decision tree model using percent U) are also plotted for comparison. **c**, Plot of the top 10 most important predictor variables, ranked by mean(| SHAP value |). **d**, SHAP values against minimal free energy (mfe) demonstrates a monotonic effect of mfe on attenuation (left), and SHAP values against percent U (pU) depicts a nonlinear effect on attenuation (right). pGC is also anticorrelated with mfe, and thus positively correlated with attenuation (left), while sequences with long homopolymer Us are responsible for the attenuation effect seen with high percent Us (right).

Because our initial model included only cell-type agnostic simple sequence features, we expanded our model to incorporate additional features related to cell-type-specific miRNA and RBP occupancy sites. However, these features did not improve model accuracy (**Supplementary Fig. 5c,d**). This finding suggests that simple, cell type-agnostic sequence features explain the predominant proportion of regulatory activity in 3’UTRs, consistent with recent studies showing that RBPs tend to bind low-complexity sequence motifs (Dominguez et al., 2018; Litterman et al., 2019). Our observation that 84.3% of the tested 3’UTR elements with significant activity in MPRAu are shared across multiple cell types is concordant with this result.

From our models that utilized only sequence-specific annotations, several low-complexity features were found to be important for prediction using SHapley Additive exPlanations (SHAP) (Lundberg and Lee, 2017). These features included homopolymer length, sequence diversity, and various features related to uracil content such as percent U/UC and UA/UU dinucleotide count (**Fig. 3c**). Investigating the individual effects of some of these important features, we found minimal free energy (mfe) monotonically negatively correlated with attenuation prediction. We also found percent GC to be anticorrelated with mfe and therefore positively correlated with attenuation (**Fig. 3d**). Surprisingly, we discovered the proportion of uracil content to have a nonlinear effect on attenuation with both low and high uracil content displaying attenuating effects. Specifically, longer uracil homopolymers demonstrate the most attenuating activity (**Fig. 3d**), concordant with their function as binding motifs for many RBPs (Dominguez et al., 2018; Mukherjee et al., 2019). The identification of attenuation features may be useful for constructing synthetic 3’UTRs with precise expression levels.

### MPRAu allelic effects are reflected in gene expression and human phenotype changes

After demonstrating MPRAu’s ability to detect the activity of 3’UTR elements, we asked whether our tamVar allelic effects are supported by causal alleles altering gene expression or phenotypic traits. We compared tamVars in GM12878 to cell-type matched allele-specific expression (ASE) data from heterozygous individuals in the Geuvadis RNA-seq dataset (Lappalainen et al., 2013) and used this comparison to estimate the positive predictive value (PPV) for tamVars in the assay. We observed moderately strong concordance between our tamVars and endogenously observed ASE (66.1% directionality agreement, binomial p=0.011) (**Fig. 4a**), which corresponds to a PPV of 32%. With higher stringency ASE calls (two-sided *t*-test p<0.001, Methods), agreement in directionality increased to 77.5% (binomial p=6.8×10^−4^, PPV of 55%). We obtained a weaker concordance when overlapping with Geuvadis expression quantitative trait loci (eQTL) data (60.5% agreement in directionality, binomial p=0.22, PPV of 20.9%) (**Supplementary Fig. 6a**), potentially due to varying regulatory factors (i.e. RBP/miRNA concentrations) muting true allelic effects when aggregating across individuals.

**Figure 4:**
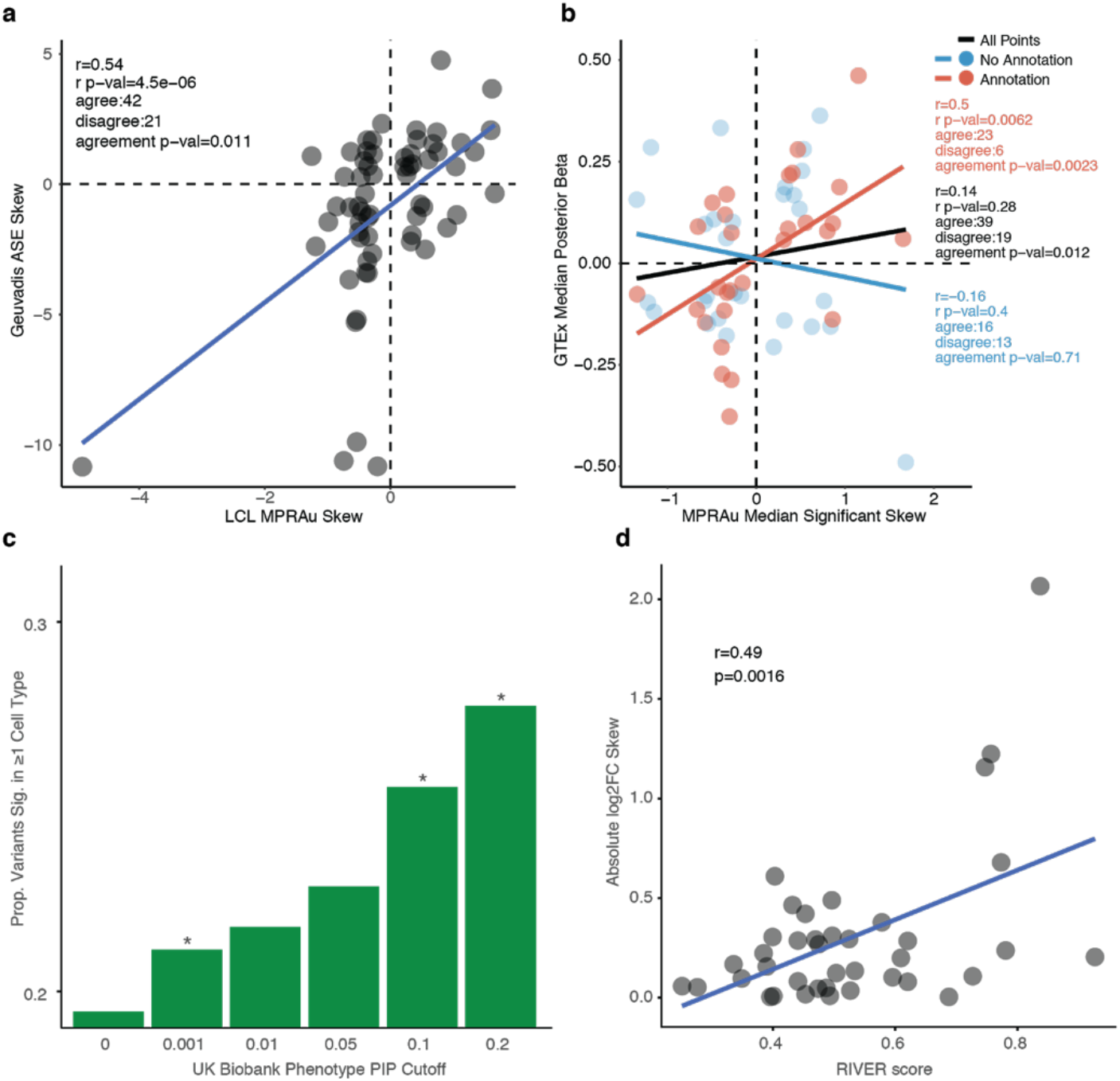
tamVars are responsible for gene expression and phenotype changes. **a**, GM12878 tamVar skew is correlated with cell-type-matched Geuvadis ASE skew. **b**, Correlation of GTEx average posterior beta across all tissues with significant effects vs tamVar average skew across all cell types demonstrates strong concordance with a high PIP cutoff (>0.2). **c**, (* p-value<0.05, Fisher’s exact test) Increasing PIP cutoffs shows a greater proportion of variants significant in at least one tested cell type, demonstrating MPRAu’s ability to capture true causal variants. **d**, tamVar skews (data for HMEC is representative and shown) are positively correlated with RIVER score, a functionality estimate of rare variants.

Next, we expanded our analysis to compare tamVars with tissue eQTLs from GTEx (GTEx Consortium, 2013), where we obtained putative causal alleles from genetic fine-mapping (Benner et al., 2016; Ulirsch; Wang et al., 2020a) (**Supplementary Table 4**). Aggregating allelic effects across cell types and tissues, we observed variants with a high inferred probability of causality (posterior inclusion probability (PIP)>0.2) displayed significant agreement in directionality between aggregated MPRAu and GTEx median effect sizes (concordance=67%, binomial p=0.012) (**Fig. 4b**). Subsetting on known 3’UTR annotations improved the concordance (82%, binomial p=9.1×10^−4^). Even at a relaxed significance threshold, we still observed significant concordance (61%, binomial p=0.04), suggesting our findings are robust (**Supplementary Fig. 7a**). We demonstrate that MPRAu is highly specific to causal variants, as the agreement in directionality is lost at lower PIPs (**Supplementary Fig. 7b**). PPV estimates based on GTEx tissue-aggregated median effect sizes (34%-64%) are similar to those based on Geuvadis ASE (32%-55%).

We next looked for enrichment of MPRAu tamVars in genetically fine-mapped causal variants associated with 94 traits in the UK Biobank (Benner et al., 2016; Ulirsch; Wang et al., 2020a). We observed greater enrichment in MPRAu functionality with increasing causality (PIP) thresholds (two-sided *t*-test p=3.64×10^−4^) (**Fig. 4c, Supplementary Table 4**). This suggests that in addition to causing in vivo gene expression changes, the tamVars identified in our study have phenotypic consequences and that MPRAu is a powerful approach for dissecting association studies.

As an orthogonal approach to confirm the tamVars identified by MPRAu were correlated with in vivo expression changes, we also assayed a set of rare variants associated with large transcriptional effects (Li et al., 2017). When we compared MPRAu allelic skews with a rare variant functionality metric (RIVER score), we observed a significant positive correlation (Pearson corr=0.42, p=7.2×10^−3^) (**Fig. 4d**). This finding suggests MPRAu can identify functionality in common as well as rare 3’UTR variants, which modern association studies have a lower power to detect.

### MPRAu SNV and deletion tiling dissects functional sequence motifs

To characterize the molecular mechanisms driving the regulatory effects of several highly significant tamVars with base-pair resolution, we created a MPRAu array that tiled single-nucleotide and deletion variants surrounding the variant (3’UTR tiling). Specifically, we extended MPRAu in two ways: we assayed 1) 5 bp non-overlapping deletions over the entire tested 3’UTR sequence, and 2) all single-nucleotide changes within +/- 10 bp of the tamVar. We carried out this deep analysis on 80 tamVars (**Supplementary Table 1**), of which three are described.

We found 3’UTR tiling useful to identify RBP sequence motifs. At rs16975240 (*FAM92B*), which in our screen displayed a strong allelic skew (log_2_FC=2.06, BH p-adj=1.33×10^−60^), the ref allele demonstrated strong expression attenuation (log_2_FC=-2.29). However, the alt allele exhibited a muted effect (log_2_FC=-0.24), which suggests the perturbation of an attenuating element. Deletions between the variant position and 10 bps upstream on the ref background identified such an element that when removed, alleviated this attenuation (+3.55-4.46 FC increase). As expected in the alt background, deletions of the same sequence yielded minimal changes (+1.13-1.16 FC increase). Saturation mutagenesis identified a 10 bp motif in the ref region perfectly matching the U1 snRNP binding sequence consensus (**Fig. 5a**). This core component of the spliceosome has also been shown to suppress gene expression by limiting poly-A tail addition (Furth et al., 1994; Guan et al., 2007). The alt allele disrupts this motif, likely explaining the underlying allelic skew.

**Figure 5:**
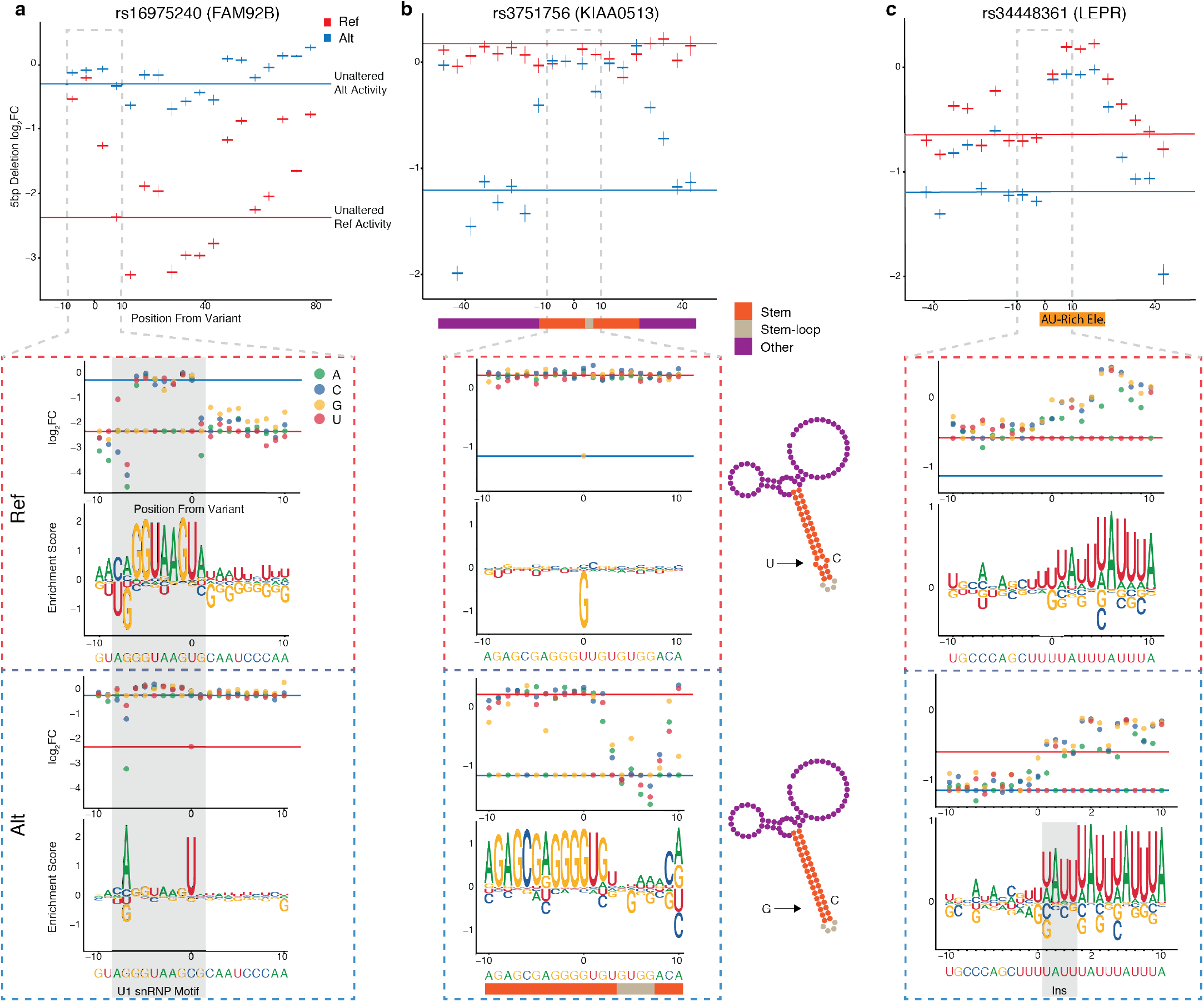
MPRAu uncovers functional sequence architectures via SNV and deletion tiling. For **a-c**, Top, 5bp deletion tiling of the targeted variant using the ref (red) or alt (blue) sequence context across 100 bp. Bottom, SNV tiling showing log2 fold change and motif enrichment score in both sequence contexts. HEK293 3’UTR activity data is plotted for all panels. **a**, Plots for tiling at rs16975420 (*FAM92B*), showing the disruption of the sequence motif for U1 snRNP (gray box). **b**, Plots for tiling at rs3751756 (*KIAA0513*), showing the disruption of a highly structured RNA arrangement (right). SNV tiling demonstrates that the greatest functional change is via disruption of bases in the predicted stem structure. **c**, Plots for tiling at rs34448361 (*LEPR*), showing the disruption of the AU-rich element.

3’UTR tiling also can be used to detect three-dimensional structures that influence RNA stability. rs3751756 (*KIAA0513*), displayed a large allelic skew (log_2_FC=-1.38, BH p-adj=2.65×10^−40^), with the alt allele demonstrating a significant attenuating effect that is unobserved with the ref allele. On the alt background, deletions within a 45 bp window around the tamVar restored expression to near ref levels, which suggests an overlying functional element. RNAfold (Gruber et al., 2008) predicts a stem-loop structure in the region, potentially mediating this attenuation (**Fig. 5b**). While the non-functional ref allele (U) creates a bulge in the stem, the alt allele (G) is predicted to create a stable stem, potentially amenable to RBP occupancy and subsequent functional attenuation. Moreover, nearly every base alteration disrupting pairing of the stem in our saturation mutagenesis alleviated repression (average Z-test p=6.13×10^−4^), in contrast to tolerated changes in the loop that yielded attenuation. Highlighting the sensitivity of MPRAu to small effect sizes, we recapitulated the expected effects of wobble base pairing, observing A to G mutations along the stem have the smallest effect sizes.

We similarly investigated the indel tamVar rs34448361 (*LEPR*) due to its large allelic skew (log_2_FC=-0.55, BH p-adj=2.19×10^−11^), and its previous identification as being within a genomic region with evidence of recent natural selection (Grossman et al., 2010). Both 3’ UTR alleles displayed attenuation, but the alt allele exhibited a significantly stronger effect (ref log_2_FC=-0.56, alt log_2_FC=-1.12), which was validated in a luciferase assay (two-sided *t*-test p=4.4×10^−4^) (**Supplementary Fig. 8a**). Deletions mapped a 25 bp AU-rich element consisting of AUUUA pentamer repeats which are known to impose attenuating effects (Siegel et al., 2020) **(Fig. 5c**). The ref 3’UTR contains four pentamers while the alt allele is a 4 bp insert that creates a fifth pentamer which is in agreement with the alt allele showing an increased effect. SNV tiling recapitulated this finding, with only disruption of AUUUA pentamers abrogating attenuation in both allelic contexts (**Fig. 5c**). The alt allele is highly represented in East Asian populations (allele frequency=0.79) versus other populations (African allele frequency=0.15) (1000 Genomes Project Consortium et al., 2015). rs34448361 is in the 3’UTR of a *LEPR* isoform (*LepRb*) (**Supplementary Fig. 8b,c**), and matches the direction of effect for an eQTL for *LepRb* across diverse tissues contexts including adipose (eQTL β=-0.26, p=2.35×10^−7^) and IFN stimulated macrophage (eQTL β=-0.50, p=6.61×10^−5^) (Kerimov et al., 2020). This isoform has been implicated in traits highly essential for survival: satiety and obesity (Münzberg and Morrison, 2015), as well as activation of various immune cells (Abella et al., 2017). However, the region surrounding *LEPR* has also been implicated in cold adaptation in modern and archaic humans (Sazzini et al., 2014). While the exact phenotype under selection is still unknown, MPRAu provides strong functional evidence for rs34448361 as a causal variant driving the evolutionary signature in the genomic region and uncovers the variant’s regulatory mechanism.

### MPRAu identifies causal 3’UTR variants related to human evolution and disease

A key attribute of MPRAu is the ability to parse genetic associations based on functional evidence to nominate causal variants for a variety of traits and diseases. Amongst all 2,153 GWAS-associated 3’UTR variants tested, we found 677 GWAS loci with a significant tamVar in at least one cell type. From this tamVar set, several previously nominated causal variants were identified, including rs13702, which disrupts a miRNA binding site in *LPL* and has been suggested to underlie associations for triglyceride levels and type 2 diabetes (Ban et al., 2010; Richardson et al., 2013; Tang et al., 2010). Our set also provides novel functional evidence for 10 variants convincingly nominated for causality from genetics alone, such as rs5891007 (*LSM1*, Schizophrenia risk) (Shi et al., 2011) and rs1140711 (*LIN7C*, bone mineral density) (Kemp et al., 2014), among others (**Table 1**). While MPRAu nominates hundreds of phenotypically-relevant 3’UTR variants, a potential limitation is the episomal-based assay not capturing endogenous gene expression effects. To more directly confirm the effects of tamVars, we performed endogenous allelic replacements via CRISPR-mediated homology-directed repair (HDR) on two tamVars linked to specific human disease processes: rs705866 and rs1059273.

rs705866 (*PILRB*) is associated with age-related macular degeneration (odds ratio (OR)=1.13, p=5×10^−9^), residing amongst 151 SNPs in strong LD to a tag-SNP (r^2^=0.808 with rs7803454) (Fritsche et al., 2016) **(Fig. 6a**). MPRAu identified a suggestive allelic effect (log_2_FC Skew=0.3, BH p-adj=0.08) with the ref allele showing an attenuating effect on expression and the alt allele disrupting a predicted miRNA binding site (hsa-miR-374a-5p) (**Fig. 6b**). Other variants in this credible set (the smallest set of variants that contain the true causal variant with 95% probability) include multiple missense SNPs, although their impacts are predicted to be benign (**Supplementary Fig. 9a,b**). Recent work identified rs705866 as an eQTL for *PILRB* specifically within the focal disease tissue, the retinal macular region (eQTL β=1.25, p=5.72×10^−29^) (Orozco et al., 2020).

**Figure 6:**
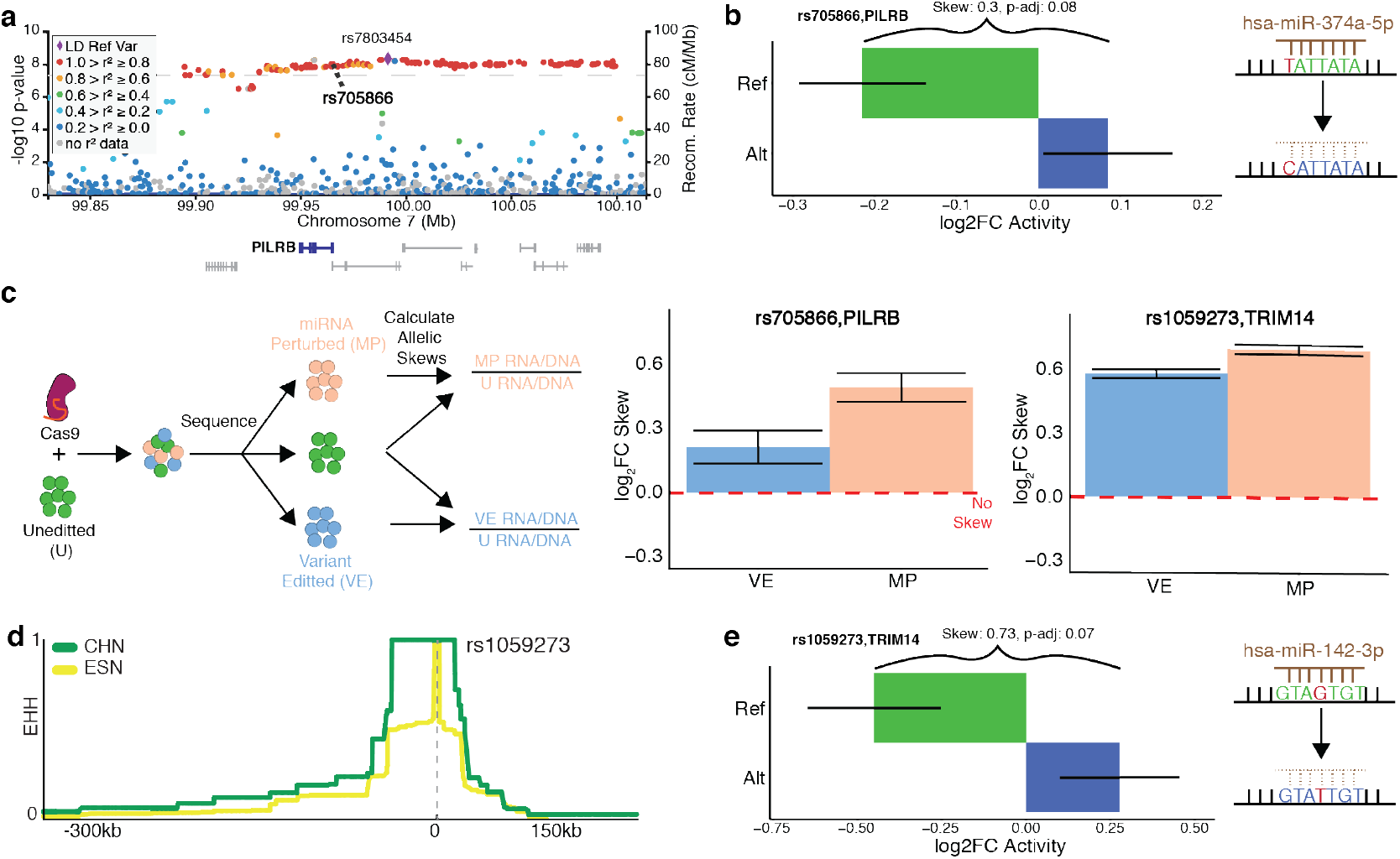
rs7803454 and rs705866 are endogenously validated tamVars impacting human disease and adaptation phenotypes. **a**, Age-related macular degeneration GWAS association plot surrounding the tag SNP rs7803454 with rs705866 (MPRAu tested SNP) in bold. **b**, MPRAu shows significant attenuating activity in the allelic background (ref) with the unperturbed miRNA binding site for rs705866. **c**, Schematic of the HDR experimental pipeline (left), HDR allelic skew results for rs705866 (center), and rs1059273 (right) recapitulate MPRAu results. **d**, EHH score surrounding rs1059273 suggests evolutionary selection in Han Chinese (CHN). For comparison, EHH scores for Esan in Nigeria (ESN) are also shown. **e**, MPRAu shows significant attenuating activity in the allelic background (ref) with the unperturbed miRNA binding site for rs1059273.

To confirm the MPRAu allelic effect in the endogenous genomic context, we used CRISPR HDR to perform allelic replacement of rs705866 (T (ref) to C (alt)) in neuronal SK-N-SH cells. Allelic ratios of RNA and DNA from cells with the desired edit were compared with unperturbed cells, and an allelic skew was found (log_2_FC=0.21, Fisher p=3.39×10^−8^) to be concordant with the effects measured by MPRAu (**Fig. 6c**). In addition, due to the imperfect editing from CRISPR HDR, a large proportion of cells also contained unspecific SNVs and indels edits over the miRNA binding site. Aggregating the effects of these imprecise edits led to an even larger functional disruption (log_2_FC=0.49, Fisher p=1.90×10^−48^) (**Fig. 6c**), providing further support for rs705866’s function via hsa-miR-374a-5p.

rs1059273 (*TRIM14*) lies within a genomic region that experienced positive selection in East Asian populations (Grossman et al., 2010). Han Chinese genomes display an extended haplotype heterozygosity score surrounding rs1059273, but identifying a causal allele has remained difficult (**Fig. 6d**). MPRAu identified attenuating effects on the ref background, but not the alt (log_2_FC Skew=0.73, BH p-adj=0.07) (**Fig. 6e**). rs1059273 disrupts a miRNA binding site (hsa-miR-142-3p), potentially explaining higher expression from the alt allele (**Fig. 6e**). *TRIM14* has many functions, including antiviral and antimicrobial activity as part of the Type I Interferon pathway (Chen et al., 2016; Tan et al., 2017; Zhou et al., 2014). Knockouts of *TRIM14* in macrophages more effectively control Mycobacterium tuberculosis replication (Hoffpauir et al., 2020), and *TRIM14* has also been found to suppress Influenza A replication (Wu et al., 2019). rs1059273 is a significant eQTL for *TRIM14* in T cells and NK cells (T cell, CD4, Th1/17, eQTL β=0.91, p=7.2×10^−9^; NK cell, CD56dim CD16+, eQTL β=0.90, p=9.7×10^−9^) (Schmiedel et al., 2018). Fine-mapping also assigns a high likelihood of the variant causally affecting *TRIM14* expression (PIP=0.53, 0.55 in Th1, Th17 cells), and finds no other potentially causal (PIP≥0.1) variants with functional annotations within the credible set containing rs1059273 (Kerimov et al., 2020). These associations further link this variant’s potential evolutionary impact to an immunological role.

We performed allelic replacement of rs1059273 (G (ref) to T (alt)) in lymphoblastoid cells and confirmed the variant’s effects on *TRIM14* (allele replacement: log_2_FC=0.58, Fisher p=6.79×10^−699^). Similar to rs708566, we aggregated the effects from cells with unspecific edits over the miRNA binding site mutations and observed an effect size larger than by rs1059273 alone (log_2_FC=0.69, Fisher p=3.32×10^−953^) (**Fig. 6c**).

## Discussion

We developed MPRAu, a high-throughput tool to functionally characterize 3’UTR variants and used it to identify 2,368 3’UTR variants that modulate transcript abundance across six cell lines. We built powerful predictive models of 3’UTR function and identified novel modes of 3’UTR regulation. This resource characterizing GWAS, selection signals, and common variation in 3’UTRs will be useful to ongoing future studies of human adaptation and disease. We expect MPRAu will be a common experimental paradigm to test variants of unknown significance and rare variants going forward. In the future, MPRAu may be further modified to specifically detect variants impacting a particular regulatory mechanism of interest, such as transcription termination (Shalem et al., 2015) or mRNA localization (Andreassi and Riccio, 2009; Berkovits and Mayr, 2015; Tushev et al., 2018).

While MPRAu bridges a tremendous gap linking 3’UTR genomic variation, functional effects, and ultimately phenotypes, the assay has important limitations. The assay’s episomal-based nature, and the limited (100 bp) variant sequence context tested, may prevent the full recapitulation of the endogenous expression effects. Furthermore, RNA steady-state levels may not be perfectly linked with protein levels (Battle et al., 2015; Chick et al., 2016); however, flow-cytometry assays have shown overwhelmingly strong agreement of RNA expression from episomal reporter assays with protein abundance (Oikonomou et al., 2014; Zhao et al., 2014).

In addition to our reporter assay, we provide additional evidence using 3’UTR tiling and endogenous allelic replacement for three variants (rs1059273, rs705866, and rs34448361) with potentially important consequences to understanding human disease and evolution. Further experiments are needed to assess each candidate variant’s contribution in the relevant cellular context. For example, while *TRIM14* expression is suppressed in multiple viral infections, including the SARS-CoV-2 virus responsible for the COVID-19 pandemic (Blanco-Melo et al., 2020), the exact extent of rs1059273 in modulating viral infectivity is unknown. While we tested our assay across six different cellular contexts, additional phenotypic-relevant variants may be found when applying our assay in disease-specific tissues.

Currently, potentially causal variants in 3’UTR elements underlying complex human diseases are largely overlooked because of the lack of tools to characterize them. The vast addition of 3’UTR measurements, especially in the context of phenotype-relevant genetic variation, may additionally inform future models of genome function. In total, MPRAu provides a framework for prioritizing regulatory variation in 3’UTRs based on functionality. Our study helps further a more comprehensive understanding of the regulatory processes important for non-coding variant function.

## Acknowledgments

We thank S. Schaffner, S. Gosai, S. Weingarten-Gabbay, A.E. Lin, D. Kotliar, C. Myhrvold, and C. Tomkins-Tinch for thoughtful conversations and help with editing the manuscript. We thank D.P. Bartel as well as his lab for thoughtful discussions. We thank I. Shlyakhter for providing the CMS regions. This work, D. Griesemer, J.R. Xue, S.K. Reilly and R. Tewhey were supported as an ENCODE Functional Characterization Center (UM1HG009435), a Broad SPARC grant, NSF DEB-1401237, and the Howard Hughes Medical Institute. D. Griesemer was partially supported by F30GM114940. S.K. Reilly was partially supported by K99HG010669 and F32HG00922. R. Tewhey is supported by R00HG008179.

## Author Contributions

D.G., R.T., and P.C.S. conceived and began the study. J.R.X. and S.K.R. performed the main analyses and completed the study. D.G., J.R.X., S.K.R., and K.K. performed experiments. J.U. and M.K. provided the fine-mapping datasets. J.D. and S.B.M. provided the rare variant dataset. C.D.N. provided help with polysome-profiling experiments. J.U., M.K., and D.K.Y. provided additional computational insights. S.K.R., R.T., and P.C.S. supervised the study.

## References

1000 Genomes Project Consortium, Auton, A., Brooks, L.D., Durbin, R.M., Garrison, E.P., Kang, H.M., Korbel, J.O., Marchini, J.L., McCarthy, S., McVean, G.A., et al. (2015). A global reference for human genetic variation. Nature 526, 68–74.

Abella, V., Scotece, M., Conde, J., Pino, J., Gonzalez-Gay, M.A., Gómez-Reino, J.J., Mera, A., Lago, F., Gómez, R., and Gualillo, O. (2017). Leptin in the interplay of inflammation, metabolism and immune system disorders. Nat. Rev. Rheumatol. 13, 100–109.

Andreassi, C., and Riccio, A. (2009). To localize or not to localize: mRNA fate is in 3′UTR ends. Trends Cell Biol. 19, 465–474.

van Arensbergen, J., Pagie, L., FitzPatrick, V.D., de Haas, M., Baltissen, M.P., Comoglio, F., van der Weide, R.H., Teunissen, H., Võsa, U., Franke, L., et al. (2019). High-throughput identification of human SNPs affecting regulatory element activity. Nat. Genet. 51, 1160–1169.

Bakheet, T., Frevel, M., Williams, B.R., Greer, W., and Khabar, K.S. (2001). ARED: human AU-rich element-containing mRNA database reveals an unexpectedly diverse functional repertoire of encoded proteins. Nucleic Acids Res. 29, 246–254.

Ban, H.-J., Heo, J.Y., Oh, K.-S., and Park, K.-J. (2010). Identification of Type 2 Diabetes-associated combination of SNPs using Support Vector Machine. BMC Genet. 11, 26.

Battle, A., Khan, Z., Wang, S.H., Mitrano, A., Ford, M.J., Pritchard, J.K., and Gilad, Y. (2015). Impact of regulatory variation from RNA to protein. Science 347, 664–667.

Benner, C., Spencer, C.C.A., Havulinna, A.S., Salomaa, V., Ripatti, S., and Pirinen, M. (2016). FINEMAP: efficient variable selection using summary data from genome-wide association studies. Bioinformatics 32, 1493–1501.

Berkovits, B.D., and Mayr, C. (2015). Alternative 3′ UTRs act as scaffolds to regulate membrane protein localization. Nature 522, 363–367.

Blanco-Melo, D., Nilsson-Payant, B.E., Liu, W.-C., Uhl, S., Hoagland, D., Møller, R., Jordan, T.X., Oishi, K., Panis, M., Sachs, D., et al. (2020). Imbalanced Host Response to SARS-CoV-2 Drives Development of COVID-19. Cell 181, 1036-1045.e9.

Bogard, N., Linder, J., Rosenberg, A.B., and Seelig, G. (2019). A Deep Neural Network for Predicting and Engineering Alternative Polyadenylation. Cell 178, 91-106.e23.

Buniello, A., MacArthur, J.A.L., Cerezo, M., Harris, L.W., Hayhurst, J., Malangone, C., McMahon, A., Morales, J., Mountjoy, E., Sollis, E., et al. (2019). The NHGRI-EBI GWAS Catalog of published genome-wide association studies, targeted arrays and summary statistics 2019. Nucleic Acids Res. 47, D1005–D1012.

Chen, M., Meng, Q., Qin, Y., Liang, P., Tan, P., He, L., Zhou, Y., Chen, Y., Huang, J., Wang, R.-F., et al. (2016). TRIM14 Inhibits cGAS Degradation Mediated by Selective Autophagy Receptor p62 to Promote Innate Immune Responses. Mol. Cell 64, 105–119.

Chick, J.M., Munger, S.C., Simecek, P., Huttlin, E.L., Choi, K., Gatti, D.M., Raghupathy, N., Svenson, K.L., Churchill, G.A., and Gygi, S.P. (2016). Defining the consequences of genetic variation on a proteome-wide scale. Nature 534, 500–505.

Choi, J., Zhang, T., Vu, A., Ablain, J., Makowski, M.M., Colli, L.M., Xu, M., Hennessey, R.C., Yin, J., Rothschild, H., et al. (2020). Massively parallel reporter assays of melanoma risk variants identify MX2 as a gene promoting melanoma. Nat. Commun. 11, 2718.

Dey, K.K., Xie, D., and Stephens, M. (2018). A new sequence logo plot to highlight enrichment and depletion. BMC Bioinformatics 19, 473.

Dominguez, D., Freese, P., Alexis, M.S., Su, A., Hochman, M., Palden, T., Bazile, C., Lambert, N.J., Van Nostrand, E.L., Pratt, G.A., et al. (2018). Sequence, Structure, and Context Preferences of Human RNA Binding Proteins. Mol. Cell 70, 854-867.e9.

Finucane, H.K., ReproGen Consortium, Schizophrenia Working Group of the Psychiatric Genomics Consortium, The RACI Consortium, Bulik-Sullivan, B., Gusev, A., Trynka, G., Reshef, Y., Loh, P.-R., Anttila, V., et al. (2015). Partitioning heritability by functional annotation using genome-wide association summary statistics. Nat. Genet. 47, 1228–1235.

Friedländer, M.R., Mackowiak, S.D., Li, N., Chen, W., and Rajewsky, N. (2012). miRDeep2 accurately identifies known and hundreds of novel microRNA genes in seven animal clades. Nucleic Acids Res. 40, 37–52.

Friedman, R.C., Farh, K.K.-H., Burge, C.B., and Bartel, D.P. (2009). Most mammalian mRNAs are conserved targets of microRNAs. Genome Res. 19, 92–105.

Fritsche, L.G., Igl, W., Bailey, J.N.C., Grassmann, F., Sengupta, S., Bragg-Gresham, J.L., Burdon, K.P., Hebbring, S.J., Wen, C., Gorski, M., et al. (2016). A large genome-wide association study of age-related macular degeneration highlights contributions of rare and common variants. Nat. Genet. 48, 134–143.

Furth, P.A., Choe, W.T., Rex, J.H., Byrne, J.C., and Baker, C.C. (1994). Sequences homologous to 5’ splice sites are required for the inhibitory activity of papillomavirus late 3’ untranslated regions. Mol. Cell. Biol. 14, 5278–5289.

Galgano, A., Forrer, M., Jaskiewicz, L., Kanitz, A., Zavolan, M., and Gerber, A.P. (2008). Comparative analysis of mRNA targets for human PUF-family proteins suggests extensive interaction with the miRNA regulatory system. PloS One 3, e3164.

Grossman, S.R., Shlyakhter, I., Shylakhter, I., Karlsson, E.K., Byrne, E.H., Morales, S., Frieden, G., Hostetter, E., Angelino, E., Garber, M., et al. (2010). A composite of multiple signals distinguishes causal variants in regions of positive selection. Science 327, 883–886.

Grossman, S.R., Andersen, K.G., Shlyakhter, I., Tabrizi, S., Winnicki, S., Yen, A., Park, D.J., Griesemer, D., Karlsson, E.K., Wong, S.H., et al. (2013). Identifying recent adaptations in large-scale genomic data. Cell 152, 703–713.

Gruber, A.R., Lorenz, R., Bernhart, S.H., Neubock, R., and Hofacker, I.L. (2008). The Vienna RNA Websuite. Nucleic Acids Res. 36, W70–W74.

GTEx Consortium (2013). The Genotype-Tissue Expression (GTEx) project. Nat. Genet. 45, 580– 585.

GTEx Consortium (2017). Genetic effects on gene expression across human tissues. Nature 550, 204–213.

Guan, F., Caratozzolo, R.M., Goraczniak, R., Ho, E.S., and Gunderson, S.I. (2007). A bipartite U1 site represses U1A expression by synergizing with PIE to inhibit nuclear polyadenylation. RNA N. Y. N 13, 2129–2140.

Gusev, A., Lee, S.H., Trynka, G., Finucane, H., Vilhjálmsson, B.J., Xu, H., Zang, C., Ripke, S., Bulik-Sullivan, B., Stahl, E., et al. (2014). Partitioning heritability of regulatory and cell-type-specific variants across 11 common diseases. Am. J. Hum. Genet. 95, 535–552.

Hafner, M., Landthaler, M., Burger, L., Khorshid, M., Hausser, J., Berninger, P., Rothballer, A., Ascano, M., Jungkamp, A.-C., Munschauer, M., et al. (2010). Transcriptome-wide identification of RNA-binding protein and microRNA target sites by PAR-CLIP. Cell 141, 129–141.

Hoffpauir, C.T., Bell, S.L., West, K.O., Jing, T., Wagner, A.R., Torres-Odio, S., Cox, J.S., West, A.P., Li, P., Patrick, K.L., et al. (2020). TRIM14 Is a Key Regulator of the Type I IFN Response during Mycobacterium tuberculosis Infection. J. Immunol. 205, 153–167.

Holcik, M., and Liebhaber, S.A. (1997). Four highly stable eukaryotic mRNAs assemble 3’ untranslated region RNA-protein complexes sharing cis and trans components. Proc. Natl. Acad. Sci. U. S. A. 94, 2410–2414.

Kemp, J.P., Medina-Gomez, C., Estrada, K., St Pourcain, B., Heppe, D.H.M., Warrington, N.M., Oei, L., Ring, S.M., Kruithof, C.J., Timpson, N.J., et al. (2014). Phenotypic Dissection of Bone Mineral Density Reveals Skeletal Site Specificity and Facilitates the Identification of Novel Loci in the Genetic Regulation of Bone Mass Attainment. PLoS Genet. 10, e1004423.

Kerimov, N., Hayhurst, J.D., Manning, J.R., Walter, P., Kolberg, L., Peikova, K., Samoviča, M., Burdett, T., Jupp, S., Parkinson, H., et al. (2020). eQTL Catalogue: a compendium of uniformly processed human gene expression and splicing QTLs (Genomics).

Kircher, M., Xiong, C., Martin, B., Schubach, M., Inoue, F., Bell, R.J.A., Costello, J.F., Shendure, J., and Ahituv, N. (2019). Saturation mutagenesis of twenty disease-associated regulatory elements at single base-pair resolution. Nat. Commun. 10, 3583.

Klein, J.C., Keith, A., Rice, S.J., Shepherd, C., Agarwal, V., Loughlin, J., and Shendure, J. (2019). Functional testing of thousands of osteoarthritis-associated variants for regulatory activity. Nat. Commun. 10, 2434.

Lappalainen, T., Sammeth, M., Friedländer, M.R., ‘t Hoen, P.A.C., Monlong, J., Rivas, M.A., Gonzàlez-Porta, M., Kurbatova, N., Griebel, T., Ferreira, P.G., et al. (2013). Transcriptome and genome sequencing uncovers functional variation in humans. Nature 501, 506–511.

Li, H. (2013). Aligning sequence reads, clone sequences and assembly contigs with BWA-MEM. ArXiv13033997 Q-Bio.

Li, X., Kim, Y., Tsang, E.K., Davis, J.R., Damani, F.N., Chiang, C., Hess, G.T., Zappala, Z., Strober, B.J., Scott, A.J., et al. (2017). The impact of rare variation on gene expression across tissues. Nature 550, 239–243.

Litterman, A.J., Kageyama, R., Le Tonqueze, O., Zhao, W., Gagnon, J.D., Goodarzi, H., Erle, D.J., and Ansel, K.M. (2019). A massively parallel 3′ UTR reporter assay reveals relationships between nucleotide content, sequence conservation, and mRNA destabilization. Genome Res. 29, 896–906.

Liu, S., Liu, Y., Zhang, Q., Wu, J., Liang, J., Yu, S., Wei, G.-H., White, K.P., and Wang, X. (2017). Systematic identification of regulatory variants associated with cancer risk. Genome Biol. 18, 194.

Love, M.I., Huber, W., and Anders, S. (2014). Moderated estimation of fold change and dispersion for RNA-seq data with DESeq2. Genome Biol. 15, 550.

Lundberg, S., and Lee, S.-I. (2017). A Unified Approach to Interpreting Model Predictions. ArXiv170507874 Cs Stat.

Magoč, T., and Salzberg, S.L. (2011). FLASH: fast length adjustment of short reads to improve genome assemblies. Bioinforma. Oxf. Engl. 27, 2957–2963.

Marbach, D., Lamparter, D., Quon, G., Kellis, M., Kutalik, Z., and Bergmann, S. (2016). Tissue-specific regulatory circuits reveal variable modular perturbations across complex diseases. Nat. Methods 13, 366–370.

Maurano, M.T., Humbert, R., Rynes, E., Thurman, R.E., Haugen, E., Wang, H., Reynolds, A.P., Sandstrom, R., Qu, H., Brody, J., et al. (2012). Systematic localization of common disease-associated variation in regulatory DNA. Science 337, 1190–1195.

McLaren, W., Gil, L., Hunt, S.E., Riat, H.S., Ritchie, G.R.S., Thormann, A., Flicek, P., and Cunningham, F. (2016). The Ensembl Variant Effect Predictor. Genome Biol. 17, 122.

Miller, C.L., Haas, U., Diaz, R., Leeper, N.J., Kundu, R.K., Patlolla, B., Assimes, T.L., Kaiser, F.J., Perisic, L., Hedin, U., et al. (2014). Coronary Heart Disease-Associated Variation in TCF21 Disrupts a miR-224 Binding Site and miRNA-Mediated Regulation. PLoS Genet. 10, e1004263.

Morris, A.R., Mukherjee, N., and Keene, J.D. (2008). Ribonomic analysis of human Pum1 reveals cistrans conservation across species despite evolution of diverse mRNA target sets. Mol. Cell. Biol. 28, 4093–4103.

Mukherjee, N., Wessels, H.-H., Lebedeva, S., Sajek, M., Ghanbari, M., Garzia, A., Munteanu, A., Yusuf, D., Farazi, T., Hoell, J.I., et al. (2019). Deciphering human ribonucleoprotein regulatory networks. Nucleic Acids Res. 47, 570–581.

Münzberg, H., and Morrison, C.D. (2015). Structure, production and signaling of leptin. Metabolism 64, 13–23.

Oikonomou, P., Goodarzi, H., and Tavazoie, S. (2014). Systematic identification of regulatory elements in conserved 3’ UTRs of human transcripts. Cell Rep. 7, 281–292.

Orenstein, Y., Wang, Y., and Berger, B. (2016). RCK: accurate and efficient inference of sequence- and structure-based protein–RNA binding models from RNAcompete data. Bioinformatics 32, i351–i359.

Orozco, L.D., Chen, H.-H., Cox, C., Katschke, K.J., Arceo, R., Espiritu, C., Caplazi, P., Nghiem, S.S., Chen, Y.-J., Modrusan, Z., et al. (2020). Integration of eQTL and a Single-Cell Atlas in the Human Eye Identifies Causal Genes for Age-Related Macular Degeneration. Cell Rep. 30, 1246-1259.e6.

Parker, S.C.J., Stitzel, M.L., Taylor, D.L., Orozco, J.M., Erdos, M.R., Akiyama, J.A., van Bueren, K.L., Chines, P.S., Narisu, N., NISC Comparative Sequencing Program, et al. (2013). Chromatin stretch enhancer states drive cell-specific gene regulation and harbor human disease risk variants. Proc. Natl. Acad. Sci. 110, 17921–17926.

Pembleton, L.W., Cogan, N.O.I., and Forster, J.W. (2013). StAMPP: an R package for calculation of genetic differentiation and structure of mixed-ploidy level populations. Mol. Ecol. Resour. 13, 946–952.

Richardson, K., Nettleton, J.A., Rotllan, N., Tanaka, T., Smith, C.E., Lai, C.-Q., Parnell, L.D., Lee, Y.-C., Lahti, J., Lemaitre, R.N., et al. (2013). Gain-of-Function Lipoprotein Lipase Variant rs13702 Modulates Lipid Traits through Disruption of a MicroRNA-410 Seed Site. Am. J. Hum. Genet. 92, 5–14.

Sample, P.J., Wang, B., Reid, D.W., Presnyak, V., McFadyen, I.J., Morris, D.R., and Seelig, G. (2019). Human 5′ UTR design and variant effect prediction from a massively parallel translation assay. Nat. Biotechnol. 37, 803–809.

Sazzini, M., Schiavo, G., De Fanti, S., Martelli, P.L., Casadio, R., and Luiselli, D. (2014). Searching for signatures of cold adaptations in modern and archaic humans: hints from the brown adipose tissue genes. Heredity 113, 259–267.

Schmiedel, B.J., Singh, D., Madrigal, A., Valdovino-Gonzalez, A.G., White, B.M., Zapardiel-Gonzalo, J., Ha, B., Altay, G., Greenbaum, J.A., McVicker, G., et al. (2018). Impact of Genetic Polymorphisms on Human Immune Cell Gene Expression. Cell 175, 1701-1715.e16.

Shalem, O., Sharon, E., Lubliner, S., Regev, I., Lotan-Pompan, M., Yakhini, Z., and Segal, E. (2015). Systematic dissection of the sequence determinants of gene 3’ end mediated expression control. PLoS Genet. 11, e1005147.

Shi, Y., Li, Z., Xu, Q., Wang, T., Li, T., Shen, J., Zhang, F., Chen, J., Zhou, G., Ji, W., et al. (2011). Common variants on 8p12 and 1q24.2 confer risk of schizophrenia. Nat. Genet. 43, 1224– 1227.

Siegel, D.A., Tonqueze, O.L., Biton, A., Zaitlen, N., and Erle, D.J. (2020). Massively Parallel Analysis of Human 3′ UTRs Reveals that AU-Rich Element Length and Registration Predict mRNA Destabilization (Genomics).

Steri, M., Orrù, V., Idda, M.L., Pitzalis, M., Pala, M., Zara, I., Sidore, C., Faà, V., Floris, M., Deiana, M., et al. (2017). Overexpression of the Cytokine BAFF and Autoimmunity Risk. N. Engl. J. Med. 376, 1615–1626.

Tan, P., He, L., Cui, J., Qian, C., Cao, X., Lin, M., Zhu, Q., Li, Y., Xing, C., Yu, X., et al. (2017). Assembly of the WHIP-TRIM14-PPP6C Mitochondrial Complex Promotes RIG-I-Mediated Antiviral Signaling. Mol. Cell 68, 293-307.e5.

Tang, W., Apostol, G., Schreiner, P.J., Jacobs, D.R., Boerwinkle, E., and Fornage, M. (2010). Associations of lipoprotein lipase gene polymorphisms with longitudinal plasma lipid trends in young adults: The Coronary Artery Risk Development in Young Adults (CARDIA) study. Circ. Cardiovasc. Genet. 3, 179–186.

Tewhey, R., Kotliar, D., Park, D.S., Liu, B., Winnicki, S., Reilly, S.K., Andersen, K.G., Mikkelsen, T.S., Lander, E.S., Schaffner, S.F., et al. (2016). Direct Identification of Hundreds of Expression-Modulating Variants using a Multiplexed Reporter Assay. Cell 165, 1519–1529.

The GTEx Consortium (2020). The GTEx Consortium atlas of genetic regulatory effects across human tissues. Science 369, 1318–1330.

Tushev, G., Glock, C., Heumüller, M., Biever, A., Jovanovic, M., and Schuman, E.M. (2018). Alternative 3′ UTRs Modify the Localization, Regulatory Potential, Stability, and Plasticity of mRNAs in Neuronal Compartments. Neuron 98, 495-511.e6.

Ulirsch, J. in prep.

Ulirsch, J.C., Nandakumar, S.K., Wang, L., Giani, F.C., Zhang, X., Rogov, P., Melnikov, A., McDonel, P., Do, R., Mikkelsen, T.S., et al. (2016). Systematic Functional Dissection of Common Genetic Variation Affecting Red Blood Cell Traits. Cell 165, 1530–1545.

Vainberg Slutskin, I., Weingarten-Gabbay, S., Nir, R., Weinberger, A., and Segal, E. (2018). Unraveling the determinants of microRNA mediated regulation using a massively parallel reporter assay. Nat. Commun. 9.

Vainberg Slutskin, I., Weinberger, A., and Segal, E. (2019). Sequence determinants of polyadenylation-mediated regulation. Genome Res. 29, 1635–1647.

Van Nostrand, E.L., Pratt, G.A., Shishkin, A.A., Gelboin-Burkhart, C., Fang, M.Y., Sundararaman, B., Blue, S.M., Nguyen, T.B., Surka, C., Elkins, K., et al. (2016). Robust transcriptome-wide discovery of RNA-binding protein binding sites with enhanced CLIP (eCLIP). Nat. Methods 13, 508–514.

Vlasova, I.A., Tahoe, N.M., Fan, D., Larsson, O., Rattenbacher, B., Sternjohn, J.R., Vasdewani, J., Karypis, G., Reilly, C.S., Bitterman, P.B., et al. (2008). Conserved GU-rich elements mediate mRNA decay by binding to CUG-binding protein 1. Mol. Cell 29, 263–270.

Wang, G., Sarkar, A., Carbonetto, P., and Stephens, M. (2020a). A simple new approach to variable selection in regression, with application to genetic fine mapping. J. R. Stat. Soc. Ser. B Stat. Methodol.

Wang, Q.S., Kelley, D.R., Ulirsch, J., Kanai, M., Sadhuka, S., Cui, R., Albors, C., Cheng, N., Okada, Y., The Biobank Japan Project, et al. (2020b). Leveraging supervised learning for functionally-informed fine-mapping of cis-eQTLs identifies an additional 20,913 putative causal eQTLs (Genomics).

Ward, L.D., and Kellis, M. (2016). HaploReg v4: systematic mining of putative causal variants, cell types, regulators and target genes for human complex traits and disease. Nucleic Acids Res. 44, D877–D881.

Welter, D., MacArthur, J., Morales, J., Burdett, T., Hall, P., Junkins, H., Klemm, A., Flicek, P., Manolio, T., Hindorff, L., et al. (2014). The NHGRI GWAS Catalog, a curated resource of SNP-trait associations. Nucleic Acids Res. 42, D1001–1006.

White, E.K., Moore-Jarrett, T., and Ruley, H.E. (2001). PUM2, a novel murine puf protein, and its consensus RNA-binding site. RNA N. Y. N 7, 1855–1866.

Wu, X., Wang, J., Wang, S., Wu, F., Chen, Z., Li, C., Cheng, G., and Qin, F.X.-F. (2019). Inhibition of Influenza A Virus Replication by TRIM14 via Its Multifaceted Protein–Protein Interaction With NP. Front. Microbiol. 10, 344.

Zhao, W., Pollack, J.L., Blagev, D.P., Zaitlen, N., McManus, M.T., and Erle, D.J. (2014). Massively parallel functional annotation of 3’ untranslated regions. Nat. Biotechnol. 32, 387–391.

Zhou, Z., Jia, X., Xue, Q., Dou, Z., Ma, Y., Zhao, Z., Jiang, Z., He, B., Jin, Q., and Wang, J. (2014). TRIM14 is a mitochondrial adaptor that facilitates retinoic acid-inducible gene-I-like receptor-mediated innate immune response. Proc. Natl. Acad. Sci. U. S. A. 111, E245–254.

