## Supplementary Information for "Genome-wide functional screen of 3’UTR variants uncovers causal variants for human disease and evolution"

### Methods

#### Variant selection and oligo design

We designed our entire tested 3'UTR sequence/variant set in two separate oligonucleotide libraries. One library contained predominantly variants in regions associated with recent human evolutionary adaptation derived from a previous scan of natural selection (Grossman et al., 2013). As this scan for natural selection relied on a method called the "Composite of Multiple Signals" (CMS), we refer to this UTR library and associated experiments as the "CMS array" throughout the paper. The other library contained variants derived from GWAS, and referred to as the "GWAS array." A full table of tested oligos/variants is listed in **Supplementary Table 1**, and a more thorough description of each of the library contents is described immediately following.

For the CMS array, we selected 9,325 common 3'UTR variants (SNPs and indels up to 10 bp in length, from Phase 1 of the Thousand Genomes Project with a global MAF≥5% amongst individuals from the pilot-phase Thousand Genomes populations) that fell within positively selected regions identified by the Composite of Multiple Signals test (Grossman et al., 2013). We additionally selected 415 common (MAF≥5%) 3'UTR variants at random from the genome.

For the GWAS array, we selected 2,153 common 3'UTR variants (SNPs and indels up to 10 bp from Phase 1 of the Thousand Genomes Project with a MAF≥5% amongst individuals from European populations) which were linked ($r^2$>0.8 using PLINK v1.9) with a variant that met genome-wide significance (p-value<$5x10^{-8}$) in the 2/13/2017 freeze of the NHGRI-EBI GWAS catalog (https://www.ebi.ac.uk/gwas). 95 indels (4.2% of the initial set of 2,248 variants) were excluded from subsequent analysis due to a reference allele coding error. We additionally included all variants from an initial pilot CMS array performed in HEK293 cells (see MPRAu transfection experiments) which significantly impacted expression (134 variants) to measure the reproducibility of variant effects across different libraries, and noticed concordant effects between the GWAS and CMS array (r=0.79, p=$2.5x10^{-28}$). Furthermore, for our 3'UTR "tiling" experiment, we incorporated SNV and deletion mutations of both the reference and alternate backgrounds of 80 variants with magnitude skew 1.5-fold or more in the CMS array (**Supplementary Table 1**). For SNV tiling, we included all SNVs from 10 bp upstream of the variant site to 10 bp downstream of the variant site. For deletion tiling, we performed 5 bp non-overlapping deletions of the entire tested 100 bp sequence surrounding the variant. Finally, we included 281 rare 3'UTR variants, 46 of which were associated with outlier expression in GTEx (Li et al., 2017) and did not have rare structural variants within 200 kb (**Supplementary Table 1**). The remaining 235 control variants were selected by first identifying non-outlier genes ((gene, individual) tuples with |Median Z-score| < 1), then extracting rare variants within 10 kb of the gene. (gene, individual) tuples with at least one rare SV within 200 kb of the gene, including the gene body, were excluded in selecting control variants. Control variants were also required to have the same gene and variant type (indel, SNP) as the outlier variants - each control variant was matched to at least one outlier variant at the same gene with the same variant type. Both rare and control variants are also annotated by the Ensembl Variant Effect Predictor (McLaren et al., 2016) as 3_prime_UTR_variant.

Both CMS and GWAS libraries were synthesized as 133 bp sequences containing a maximum of 101 bp of 3'UTR context and sequence adapters on either end (5'CGAGCTCGCTAGCCT [maximum of 101 bp of 3'UTR sequence] AGATCGGAAGAGCGTCG3') (**Fig. 1a**). To select 3'UTRs, we searched the Gencode v19 database for 3'UTR-annotated entries downstream of CDS-annotated entries, filtering out entries annotated for nonsense-mediated decay and non-stop decay. Ideally, 50 bp of 3'UTR sequence context on each side of the variant was obtained; if the variant was within 50 bp of the 5' (stop codon) or 3' (termination site) end of the 3'UTR, its sequence context ended prematurely, and if possible, additional sequence was taken from the other end to obtain 100 bp total. If a variant's oligo sequence context overlapped one or more of the other variants tested in our set, we designed a reference and alternate oligo in which all other variants were assigned the reference allele (as in the reference human genome), as well as a reference and alternate oligo in which all other variants were assigned the alternate allele. For example, a variant Y flanked by one or more variants, such as variants X and Z, would be referenced in our variant datasets within our supplementary tables as follows: Y would refer to a comparison between X_ref/Y_ref/Z_ref and X_ref/Y_alt/Z_ref, whereas Y_2 would refer to a comparison between X_alt/Y_ref/Z_alt and X_alt/Y_alt/Z_alt.

**Oligo synthesis and amplification**

The CMS array oligos were synthesized by CustomArray and the GWAS array oligos were synthesized by Twist Biosciences. Post-synthesis, for the CMS array, 6 bp random hexamer barcodes and additional adapter sequences were added by performing 20 PCR reactions, each 50 µL in volume, containing 5.7 ng of oligo, 25 µL of Q5 NEBNext MasterMix (NEB, M0541S), 1 unit Q5 HotStart polymerase (NEB, M0493S), 0.5 µM oligo_BAR_Bmt_F and oligo_pmir_R_min primers, and 20 µg BSA (NEB, B9000) **(Supplementary Table 5)**. PCR cycling conditions used are as follows: 98°C for 30 seconds, 22 cycles (98°C for 30 sec, 60°C for 30 sec, and 72°C for 1 min), 72°C for 2 min. For the GWAS array oligos, PCR was performed using the same primers and cycling conditions with the following exceptions: 1) 24 50 µL PCR reactions instead of 20, 2) 1 ng of oligo in each reaction instead of 5.7 ng, 3) use of the NEBNext Ultra II Q5 Master Mix (NEB, M0544L) instead of the Q5 HotStart polymerase, 4) and 12 cycles for amplification instead of 22 cycles. PCR reactions from both arrays were purified by performing a 2.5X SPRI purification using Agencourt AMPure XP SPRI (Beckman Coulter, A63881) beads according to manufacturer instructions.

**MPRAu vector assembly**

To create our MPRAu plasmid pool for the CMS array, barcoded oligos were inserted into a modified pmirGLO (Promega, E1330) vector, pmirGLO:Δluc::gfp $\Delta Amp^R::Kan^R$, which contains the GFP gene driven by the pgk promoter. Oligos were inserted by Gibson Assembly (NEB, E2611) using 1 µg of BmtI/XbaI (NEB, R0658S; NEB, R0145S) digested vector and 415 ng of amplified oligos (10:1 plasmid to oligo molar ratio) in a 40 µL reaction incubated for 60 min at 50°C followed by 2X SPRI purification and eluted in 25 µL EB. The elution was then concentrated to 10 µL by vacuum centrifugation. The ligated vector was then split into 5 µL aliquots, each of which was transformed into 100 µL of 10-beta e.coli (NEB, C3020K) by electroporation (2kV, 200 ohm, 25 µF). Electroporated bacteria were immediately split into five 1 mL aliquots of SOC (NEB, B9020S) and recovered for 1 hour at 37°C then independently expanded in 1L of LB supplemented with 50 µg/mL of kanamycin (Teknova, K2125) on a floor shaker at 37°C for 12 hours. After outgrowth, aliquots were pooled before plasmid purification (Qiagen, 12991). For each of the aliquots, we plated serial dilutions after SOC recovery and estimated a library size of $>10^8$ CFUs. For the GWAS array, a similar vector assembly protocol was followed except the Gibson Assembly reaction time and size was doubled. Each electroporation also was split into four 1 mL aliquots of SOC, subsequently pooled, and grown into four final 1L LB supplemental with 50 µg/mL of kanamycin in a floor shaker.

**MPRAu transfection experiments**

We performed our initial pilot experiment using the CMS array into HEK293 cells (specifically the HEK293FT cell line (Thermofisher, R70007)). HEK293 cells were grown in DMEM (ThermoFisher,

10564) supplemented with 10% FBS (ThermoFisher, A3160401), 1X NEAA (ThermoFisher, 11140050), and 1 mM Sodium Pyruvate (ThermoFisher, 11360070). For all 5 replicate transfections (Figure 1C), cells were plated in a 15 cm plate and grown to a density of ~80-90%. Cells were then transfected with 80 µL Lipofectamine 2000 (ThermoFisher, 11668027) and 20 µg DNA, and incubated with the transfection reagents for 24 hours, monitored by fluorescent microscopy. Afterward, transfected cells were then split 1:3 into 2 new 15 cm plates. After an additional 24 hours (48 hours post-transfection), media from each plate was replaced with 30 mL DMEM + 100 µg/mL cycloheximide (CHX) (Sigma-Aldrich, C4859) and incubated for 5 minutes at 37°C. Both plates were then washed with 10 mL cold PBS (ThermoFisher, 14040) + 100 µg/mL CHX, scraped in 1 mL cold PBS + 100 µg/mL CHX, pooled and centrifuged for 5 minutes at 500 x g, and finally resuspended in 350 µL lysis buffer composed of 5 mM Tris pH 7.5 (ThermoFisher, 15567), 2.5 mM Magnesium Chloride (ThermoFisher, AM9530G), 1.5 mM Potassium Chloride (ThermoFisher, AM9640G), 2 µM DTT (VWR, 97061-340), 100 µg/mL CHX, 5 mg/mL Triton-X (Sigma-Aldrich, T8787), and 5 mg/mL sodium deoxycholate (Sigma-Aldrich, 30970). The cell lysate was then centrifuged for 5 minutes at 12,000 x g and the supernatant was flash-frozen in vapor phase nitrogen. For polysome profiling, approximately 200 µL lysate was loaded onto 10%-50% sucrose gradients in 15 mM Tris pH 7.5, 15 mM Magnesium Chloride, 150mM Sodium Chloride (ThermoFisher, AM9760G), and 100 µg/mL CHX. Gradients were centrifuged in SW41Ti rotor at 35,000 rpm for 2.5 hours at 4°C and 0.5 mL fractions were collected.

We then transfected the GWAS array into HEK293 cells, following the same protocol above, except using two 15 cm plates per replicate, and excluding the polysome profiling step. The CMS & GWAS array were then pooled together into one library (CMS+GWAS) for subsequent transfections across other cell types, with protocols described below.

GM12878s (Coriell) were grown in RPMI (ThermoFisher, 61870036) supplemented with 15% FBS and 1% 10X Penicillin-Streptomycin (Pen-Strep; ThermoFisher, 15140122; Corning, 30-002-CI). Each replicate (4 total, grown on different days) was grown to ~ $1x10^6$ cells/mL, 150 million cells were then collected via centrifugation at 300 x g and suspended in 1.2 mL RPMI with 150 µg library. Subsequently, cells were electroporated with the Neon transfection system with the 100 µL kit (ThermoFisher, MPK10096) using 3 pulses of 1200V, 20 ms each. Following transfection, each replicate was grown in 150mL RPMI + 15% FBS without Pen-Strep, to recover for 48 hours. Cells were split 1:2 after the first 24 hour recovery time to prevent overgrowth. The cells were then spun down, washed once with PBS, flash-frozen via liquid nitrogen, and subsequently stored at -80°C.

K562s (ATCC, CCL-243) were grown in RPMI supplemented with 10% FBS and 1% 10X Pen-Strep. Each replicate (4 total, grown on different days) was grown to ~ $1x10^6$ cells/mL, 150 million cells were then collected via centrifugation at 300 x g and suspended in 1.2 mL RPMI with 150 µg library. Subsequently, cells were electroporated with the Neon transfection system with the 100 µL kit using 3 pulses of 1450 V, 10 ms each. Following transfection, each replicate was grown in 150 mL RPMI + 15% FBS without Pen-Strep to recover for 48 hours. Cells were split 1:2 after the first 24 hour recovery time to prevent overgrowth. The cells were then spun down, washed once with PBS, flash-frozen via liquid nitrogen, and subsequently stored at -80°C.

HepG2s (ATCC, HB-8065) were grown in 25 mL MEM Alpha (ThermoFisher, 32561037) + 10% FBS + 1% Pen-Strep on 15 cm plates. Cells were grown to 60%-80% confluency. For all 4 replicates, grown on different days, two 15 cm plates per replicate were transfected with 87.5 µL Lipofectamine 3000 (ThermoFisher, L3000015) and 35 µg library. Following transfection, each replicate was grown in 25 mL MEM Alpha + 10% FBS without Pen-Strep to recover for 48 hours. The cells were then trypsinized, spun down at 300 x g at 4°C, washed once with PBS, flash-frozen via liquid nitrogen, and subsequently stored at -80°C.

SK-N-SH (ATCC, HTB-11) were grown in 90 mL EMEM (ATCC, 30-2003) supplemented with 10% FBS and 1% Pen-Strep on Nunc Triple Flasks (VWR, 89498-706). Each replicate (4 total, grown on different days) was grown to 80%-100% confluency. Cells were trypsinized, and 40 million cells were collected and resuspended in 400 µL Buffer R with 25 µg library. Subsequently, cells were electroporated with the Neon transfection system with the 100 µL kit using 3 pulses of 950 V, 30 ms each. Following transfection, each replicate was grown in 45 mL EMEM + 10% FBS without Pen-Strep to recover for 48 hours. The cells were then trypsinized, spun down at 300 x g at 4°C, washed once with PBS, flash-frozen via liquid nitrogen, and subsequently stored at -80°C.

HMECs (ThermoFisher, A10565) were grown in 60 mL MEGM (Lonza, CC-3150) in T225 flasks. For each replicate (5 total, grown on different days), 6 confluent flasks were grown to 80%-100%. Cells were then resuspended in buffer R and DNA to get a final concentration of 10 million cells/mL and 25 µg DNA/mL. Subsequently, cells were electroporated with the Neon transfection system with the 100 µL kit using 3 pulses of 950 V, 30 ms each. Following transfection, each replicate was grown in 4 T225 flasks (each with 60 mL MEGM) to recover for 48 hours. The cells were then trypsinized, spun down at 250 x g at 4°C, washed once with PBS, flash-frozen via liquid nitrogen, and subsequently stored at -80°C.

Transfection efficiency was monitored across all cell types by assessing GFP fluorescence. Across all cell lines, greater than 50% of live cells fluoresced after transfection, signifying acquisition of the MPRA construct. K562, HepG2, and HEK293 had the highest transfection efficiency (>80%), while HMEC had the worst (50%).

**RNA extraction and cDNA synthesis**

For all cell type replicates, RNA was extracted with TRIzol LS (ThermoFisher, 10296) according to manufacturer instructions. Total RNA was purified from the cell lysate. Polysomal RNA was purified from sucrose fractions, pooling fractions corresponding to three or more ribosomes. 7.5 µg GlycoBlue (ThermoFisher, AM9515) was added to each sample to visualize the pellet. mRNA was purified from total RNA using oligo $d(T)_{25}$ magnetic beads (NEB, S1419S) according to the manufacturer's instructions, and eluted at 80°C. Purified total and polysomal mRNA were then subjected to Turbo DNase treatment (ThermoFisher, AM2239). The reaction was terminated in 2 mg/mL SDS (ThermoFisher, AM9822) and purified by performing a 2X SPRI purification using Agencourt RNAClean XP SPRI (Beckman Coulter, A63987) beads according to manufacturer instructions. DNase-treated mRNA for each of the tested replicates was diluted to the concentration of the lowest concentration sample, and first-strand cDNA was synthesized from concentration-normalized mRNA with SuperScript III (ThermoFisher, 18080) and a gene-specific primer 162 bp downstream of the oligo (oligo_RT_R, **Supplementary Table 5**). For the SuperScript III reaction, we used the manufacturer's recommended protocol, except by increasing the total reaction volume to 40 µL and performing the elongation step at 55°C for 80 minutes. Single-stranded cDNA was purified by performing a 2X SPRI purification using Agencourt RNAClean XP SPRI beads.

**qPCR and library construction**

cDNA concentrations from the cell type replicates were estimated via qPCR using 1 µL of cDNA sample in a 10 µL reaction that contained 5 µL Q5 NEBNext master mix, 1.7 µL SYBR Green I diluted 1:10,000 (Life Technologies, S-7567), and 0.5 µM of PE_PCR_P1 and PE_PCR_P2_BMT primers (**Supplementary Table 5**), and under the following conditions: 98°C for 30 seconds, 40 cycles (98°C for 10 sec, 65°C for 30 sec, and 72°C for 30 sec), 72°C for 5 minutes. The cDNA samples across all cell types had a cycle threshold (CT) between 7 and 15 cycles.

Samples across all tested replicates from all cellular cDNA samples were then aliquoted to achieve the same input going into the next amplification step based on the CT values derived from the previous step. Specifically, we matched all input amounts to achieve a CT of 11. The plasmid pool was also amplified in five independent PCR reactions (technical replicates), also adjusting the input amount per reaction to achieve an expected CT of 11. Samples were amplified in a 50 µL PCR reaction containing 25 µL NEBNext Ultra II Q5 Master Mix and 0.5 µM of PE_PCR_P1 and PE_PCR_P2_Bmt (**Supplementary Table 5**), and under the following conditions: 98°C for 30 seconds, 9 cycles (98°C for 10 sec, 65°C for 30 sec, 72°C for 30 sec), 72°C for 5 minutes. 2.5X AMPure SPRI purification was then performed, and another round of PCR (as above, except with 5 cycles) was performed using a set of Illumina P5 index primers and a set of Illumina P7 index primers in a 100 µL reaction. Another 2.5X AMPure SPRI purification was performed afterward. Samples were then pooled according to molar estimates from the Agilent 2200 TapeStation (using the D1000 screentape reagents (Agilent, 5067-5585)) and then subsequently sequenced using a S4 flowcell (2 x 150 bp) on a NovaSeq using the Broad Institute's walk-up sequencing service.

Sample preparation for the initial pilot CMS array pilot library in HEK293 cells was processed separately in an analogous manner just described, with the main difference being the use of the SYBR Green Master Mix (ThermoFisher, 4367659) to quantify CT. The pilot library samples were sequenced using 2 x 150 bp chemistry on an Illumina HiSeq through the Broad Institute's walk-up sequencing service.

**Read alignment to 3'UTR sequences and generation of 3'UTR element count table**

Paired-end 150 bp reads were merged into single amplicons using Flash v1.2.11 (flags: -M 150, -O) (Magoč and Salzberg, 2011). Amplicon sequences were retained for quantification if the sequence surrounding the barcode met the following conditions: (1) a perfect match was found to the 10 bp sequence on either the left or right side of the barcode, (2) the 10 bp sequences on both the left and right sides of the barcode matched with a Levenshtein distance of 3 or less, and (3) the 2 bp immediately surrounding each side of the barcode matched perfectly. Oligo sequences from the passing reads were then mapped back to the expected oligo sequences using BWA mem version 0.7.12 (flags: -M) (Li, 2013). We calculated our own alignment scores to assess the quality of the alignments. These scores were calculated as the number of matching bases divided by the expected oligo size. Reads with alignment scores of less than 0.95 were discarded. Oligo libraries were extremely complex, with an average of 70-330 unique hexamer barcodes per oligo per replicate sample, which would minimize effects from any outlier barcodes that would have functional effects. As a result, oligo reads were pooled across barcodes for oligo analyses and barcode reads were pooled across oligos for barcode analyses. On average, each oligo contained 1100-3300 reads across all tested cell type/plasmid replicates (**Supplementary Table 1**).

**Functional 3'UTR element and tamVar calling**

Oligo counts from all samples were passed into DESeq2 and a median-of-ratios method was used to normalize samples for varying sequencing depths (Love et al., 2014). Normalized read counts of each oligo were then modeled by DESeq2 as a negative binomial distribution. DESeq2 estimates variance for each NB by pooling all oligo counts across samples and fitting a trend line to model the relationship between oligo counts and observed dispersion. It then applies an empirical Bayes shrinkage by taking the observed relationship as a prior and performing a maximum a posteriori estimate of the dispersion for each oligo. The overall result is that DESeq2 can obtain an estimate for dispersion of each oligo with greatly reduced bias by pooling information from all oligos.

We first used DESeq2 to estimate oligo fold changes between our sample types (plasmid pool, total RNA, polysomal RNA) for our initial pilot HEK293 dataset with just the CMS array. To calculate total RNA expression and polysomal RNA expression, we normalized total RNA counts in the lysate and polysomal RNA counts respectively by the baseline counts in the plasmid pool (design = ~ Replicate+Sample_Type). We estimated fold changes between the reference and alternate alleles (RNAskew and POLYskew) by adding an interaction term (design = ~

Replicate+Sample_Type+Variant+Sample_Type:Variant) and using a Wald test with the Bonferroni multiple test correction. In all models, a replicate term was added to pair samples from the same transfection. We used this initial model to look at the concordance between polysomal and total RNA data (**Supplementary Figure 1**).

Upon expanding the assay to the 5 other cell types other than HEK293, we ran DESeq2 separately for each cell type to derive functional 3'UTR elements and tamVars using a revised model. The DESeq2 model was revised as follows: design = ~ Variant+Type+Variant:Type, where Type corresponds to total RNA or the plasmid pool. Wald tests were used with contrasts to derive reference and alternate specific activity (fold changes of RNA over plasmid) and the difference between alternate activity and reference activity (allelic skew). The Benjamini-Hochberg test correction was performed via DESeq2 to correct for multiple hypothesis testing. Variants with significant skew (tamVars) were designated if the adjusted p-value from any of the tested allelic backgrounds was less than 0.1. tamVars were shared between cell types X and Y if the adjusted p-value was less than 0.1 for both X and Y. The output from this DESeq2 analysis is the one reported in **Supplementary Table 1**.

### Luciferase assays

To validate the expression values obtained by MPRAu, we selected 18 oligos consisting of nine ref/alt pairs. Five of the oligos were selected as no-skew controls for having uncorrected p-values of greater than 0.01. We designed the same 101 bp sequence that was tested by MPRAu as a gBlock (IDT) and cloned each into the pmirGLO dual-luciferase reporter vector (Promega, E1330). Cells were plated in a 96 well plate and grown to a density of 80%-90%, then transfected with a mixture of 0.2 µL Lipofectamine 2000, 500 pg of the cloned dual-luciferase vector, and 49.5 ng of pGL4.23, a promoterless control vector (Promega, E8411). We performed six transfection replicates per oligo (all on the same 96-well plate). Cells were incubated with transfection reagents for 24 hours, monitored by fluorescent microscopy, and then split 1:3 into a new 96-well plate. After 24 hours (48 hours post-transfection), firefly and Renilla luminescence were read from each well using the Dual-Glo Luciferase Assay (Promega, E2920). Firefly luciferase luminescence for each well was normalized to the Renilla luciferase luminescence for the same well, and each experiment was normalized as a log-ratio value relative to the mean of a control oligo with an MPRAu RNA/DNA ratio of -0.2 (**Supplementary Table 2**).

### CRISPR allelic replacement

All crRNA and ssODN were designed and ordered via IDT (**Supplementary Table 5**). Cas9/Cpf1 reagents were also ordered from IDT. Two replicate experiments were performed for each target. rs1059273_GuideRNA_Cas9 (Cas9 crRNA) and rs1059273_ssODN (ssODN) were used for both replicates of rs1059273. For rs705866, rs705866_GuideRNA_Cpf1_1 (Cpf1 crRNA) and rs705866_ssODN_1 (ssODN) were used for the first replicate, and rs705866_GuideRNA_Cpf1_2 (Cpf1 crRNA) and rs705866_ssODN_2 (ssODN) were used for the second replicate. rs1059273 experiments were performed in GM12878 and rs705866 experiments were performed in SK-N-SH. Consistent effects were observed across both replicate experiments for both targets (**Fig. 6c**, **Supplementary Fig. 10a,b**). Cells were grown in the following media conditions: RPMI, supplemented with 15% FBS for GM12878s, and EMEM supplemented with 10% FBS for SK-N-SH. The HDR protocol used was adapted from IDT's provided one: http://sfvideo.blob.core.windows.net/sitefinity/docs/default-source/protocol/homology-directed-repair-alt-r-crispr-cas9-ultramer-oligos.pdf?sfvrsn=9750707_8

For the Cas9 HDR experiments, the following protocol was used per target. First, 0.9 µL of 200 µM Alt-R CRISPR-Cas9 target-specific crRNA, 0.9 µL of 200 µM Alt-R CRISPR-Cas9 tracrRNA (IDT, 1072533), and 1.5 µL Nuclease-Free Duplex Buffer (IDT, 1072570) were combined and heated at 95°C for 5 minutes. The crRNA:tracrRNA solution was then cooled at room temperature. 3 µL of the crRNA:tracrRNA solution was then combined with 2 µL Alt-R S.p. HiFi Cas9 Nuclease V3 (IDT, 1081059) and incubated at room temperature for 10-20 minutes to form the RNP complex. 100K cells per electroporation were washed with PBS, then resuspended in 7.69 µL of Neon Resuspension Buffer R. Next, 1.61 µL of the RNP complex, 7.69 µL of 100K cells in Neon Resuspension Buffer R, 0.3 µL of 100 µM ssODN, and 0.4 µL of Alt-R Cas9 Electroporation Enhancer (IDT, 1075916) were combined for one electroporation using the Neon transfection system with the 10 µL kit (ThermoFisher, MPK1025). Each target underwent two electroporations using set electroporation conditions (3 pulses of 1200 V, 30 ms each for GM12878s). Both electroporations were transferred to a well containing 0.4 mL of recovery media (regular media supplemented with 30 µM HDR enhancer (IDT, 1081072)) in a 24-well plate and grown for 12-24 hours. The recovery media was then changed to regular media afterward. Cells were grown and expanded until we achieved a population of at least 6-8 million cells, then ~ 6-8 million cells were extracted, washed with PBS, and flash-frozen afterward.

For the Cpf1 HDR experiments, the following protocol was used per target. First, 2.5 µL of Alt-R CRISPR-Cpf1 target-specific crRNA was combined with 2.5 µL Alt-R A.s. Cas12a (Cpf1) Ultra (IDT, 10001273) and incubated at room temperature for 10-20 minutes to form the RNP complex. Following the formation of the RNP complex, the protocol follows exactly as the Cas9 HDR protocol, except with the use of 0.3 µL of Alt-R Cpf1 Electroporation Enhancer (IDT, 1076300) (cells were also resuspended in 7.79 µL Neon Resuspension Buffer R instead of 7.69 µL to account for the 0.1 µL decrease in volume), and the use of a different electroporation setting for SK-N-SH (3 pulses of 950 V, 30 ms each ).

DNA/RNA was extracted from the frozen samples using the AllPrep DNA/RNA Mini Kit (Qiagen, 80204). Extracted RNA was subsequently DNase treated, terminated in 2 mg/mL SDS (ThermoFisher, AM9822), and purified via 2X SPRI using Agencourt RNAClean XP SPRI beads. DNase-treated RNA was then used to generate target-specific cDNA using SuperScript III and a gene-specific primer (rs1059273_R) for target rs1059273. For target rs705866, we used the same gene-specific primer but utilized SuperScript IV VILO instead (we switched enzymes due to the lower expression levels of the gene). 17 20 µL reactions with 500 ng RNA in each reaction were performed for target rs1059273 and 12 20 µL reactions with 500 ng RNA in each reaction for target rs705866. The entire 20 µL from each reaction was then directly used to amplify the target amplicon via PCR using the NEBNext Ultra II Q5 Master Mix with 0.5 µM rs1059273_F and rs1059273_R primers and the following cycling conditions: 95°C for 20 seconds, 15 cycles (95°C for 20 sec, 68°C for 20 sec, 72°C for 30 sec), 72°C for 2 minutes (**Supplementary Table 5**). rs705866 had the same cycling conditions except with using primers rs705866_F and rs705866_R and 12 instead of 15 cycles for amplification (**Supplementary Table 5**). Purified DNA was also amplified via PCR using the same target primers and subject to the same cycling conditions for each target, except with 50 individual PCR reactions for target rs1059273 and 12 individual PCR reactions for target rs1059273. For each target DNA/RNA, the individual post-PCR reactions were then pooled together, subject to a 1X AMPure SPRI purification, and concentrated via vacuum centrifugation. Another round of PCR was then performed (same cycling conditions as above, except with 8 cycles and 64°C for the annealing temperature) to attach p7 and p5 Illumina adapters with unique sample indices. The PCR products were then subject to another 2X SPRI and eluted in 30 µL. The resulting purified PCR products across all targets were then molar pooled from Agilent 2200 TapeStation quantifications (using D1000 screentape reagents) and subsequently sequenced using 2 x 150 bp chemistry on an Illumina MiSeq.

CRISPR HDR was found to be efficient across all replicates. For rs705866, 3.5% of alleles from the first replicate and 14.5% of the alleles from the second replicate obtained perfect edits. For rs1059273, 31.4% of the alleles from the first replicate and 34.3% of the alleles from the second replicate acquired perfect edits. Furthermore, for rs705866, 4.3% of alleles from the first replicate and 3.6% from the second replicate had additional sequence perturbations over the expected miRNA motif on the ref background, which allowed us to quantify the effects of other SNVs/indels overlying the functional element (**Fig. 6c**) Similarly, for rs1059273, 22.5% of alleles from the first replicate and 24.1% from the second replicate had additional perturbations over the expected miRNA motif on the ref background.

**Annotations used for enrichment analyses and modeling**

We identified AU-rich elements in multiple ways, including using the AUUUA pentamer that often occurs in multiples, the UUAUUUAWW nonamer that has been associated with rapid mRNA decay, and finally two 13 bp motifs from Bakheet et al. 2001 (Bakheet et al., 2001) (WWWUAUUUAUWWW and WWAUUUAUUUAWW, with one mismatch allowed in the flanking sequence outside of AUUUA and UUUAUUU respectively). We identified CU-rich elements using the (C/U)CCANxCCC (U/A) (C/U)yUC (C/U)CC consensus sequence that has been shown to increase mRNA stability (Holcik and Liebhaber, 1997). We identified GU-rich elements using the UGUUUGUUUGU consensus sequence that has been associated with short-lived transcripts (Vlasova et al., 2008). We identified Pumilio binding sites using the UGUANAUA motif identified through human Pumilio immunoprecipitation experiments (Galgano et al., 2008; Hafner et al., 2010; Morris et al., 2008).

To predict microRNA binding sites, we used TargetScan6 (Friedman et al., 2009) to identify 7mer and 8mer seed matches to the top 10 and top 100 most abundant microRNAs in each of the cell lines tested. To obtain the top 10 and top 100 most abundant microRNAs, we used miRDeep2 (Friedländer et al., 2012) to quantify microRNA abundance in short RNA sequencing experiments from ENCODE and the Sequence Read Archive (see below). For SK-N-SH we were unable to obtain sequencing data and instead used short RNA sequencing of the SH-SY5Y line, a neuroblast-like subline of SK-N-SH. To generate the heatmap in **Fig. 2d**, for each cell type with multiple miRNA datasets, we used the miRNA dataset that demonstrated the most significant effect when overlapping with the 8mer annotations from the dataset's top 10 most abundant miRNAs. Significance of effect was measured by the p-value derived from a *t*-test on the expression values of the oligos that contained an 8mer motif from the dataset's top 10 most abundant miRNAs (comparing against a null of zero, with an alternative hypothesis of the mean expression values being less than zero). We used the same best miRNA dataset per cell type to extract features for our functional 3'UTR element computational modeling work.

| **Cell Line** | **Archive** | **Dataset ID** |
|---|---|---|
| HEK293 | SRA | • SRR1240816<br>• SRR1240817 |
| SH-SY5Y | SRA | • SRR1304311 |
| HepG2 | SRA | • ERR738415<br>• ERR738403<br>• ERR738417 |
| HepG2 | ENCODE | • ENCFF175ZOB |
| HMEC | SRA | • DRR041459<br>• DRR041581<br>• DRR041472<br>• SRR5127224 |
| GM12878 | ENCODE | • ENCFF878BGO<br>• ENCFF440XMV<br>• ENCFF322QWE |
| K562 | ENCODE | • ENCFF119JCH<br>• ENCFF756CSN<br>• ENCFF691LGG |

The RBP dataset used in generating **Supplementary Fig. 2** was derived from Table S3 from Dominguez et al. 2018. Motifs with Stepwise_R-1 greater than 0.1 were classified as having a "strong RBP motif", and motifs with Stepwise_R-1 between 0 and 0.1 were classified as having a "weak RBP motif."

#### RBP expression data derivation

RNA expression data (for deriving RBP expression, and used for our computational prediction work) were downloaded as tsv files from the following sources:

| **Cell Line** | **Source** | **Link/ENCODE file accession** |
|---|---|---|
| HEK293 | Protein Atlas | https://www.proteinatlas.org/download/rna_celline.tsv.zip |
| HepG2 | ENCODE | ENCFF004HYK |
| HMEC | ENCODE | ENCFF380GBC |
| GM12878 | ENCODE | ENCFF599JTV |
| K562 | ENCODE | ENCFF172GIN |
| SK-N-SH | ENCODE | ENCFF389TFR |

#### Hexamer barcode analysis

We used the barcodes included in our MPRAu design to directly measure the activity of all possible 4,096 hexamer sequences across every tested cell type (**Fig. 1a**, **Supplementary Table 3**). For each cell type/plasmid replicate, counts were derived for each of the 4,096 hexamers by summing up the counts of all CMS array oligos that were tagged with the corresponding hexamer. For assembling the hexamer count table, only oligos that belonged to the CMS array were used to ensure that hexamers were derived from a diverse oligo pool. The GWAS oligo set was overrepresented by 3'UTR tiling oligos, comprising 69% of the GWAS oligo set. Since these 3'UTR tiling oligos were designed based on oligos significant effects in the CMS array, hexamers obtained from these oligos were inherently biased to have significant effects. To not bias hexamer activity with interactions with these oligos, the GWAS oligos were excluded from calculating hexamer activity. DESeq2 was also utilized to calculate the fold change of RNA over DNA independently for each cell type. The DESeq2 model used was design = ~ Type, where Type corresponds to RNA or the plasmid backbone. The Wald test was used to derive p-values, and the Benjamini-Hochberg test correction was performed via DESeq2 to correct for multiple hypothesis testing. Using the DESeq2 derived adjusted p-values, we were able to derive the hexamers having the most significant effects per cell type.

Examining the hexamers with the most significant effects ($|\log_2\text{FC Skew}| \geq 0.5$, BH p-adj<0.1), we identified up to 28 hexamers per cell type with large attenuating or augmenting effects. Our tested hexamers confirmed effects expected at known RBP motifs (Dominguez et al., 2018) (**Supplementary Fig. 2a**) and variants perturbing functional hexamers abrogated the intact element's effect (two-sided Wilcoxon rank-sum test $p<0.05$ across most tested cell types) (**Supplementary Fig. 3a**,**b**). As each 3'UTR was linked to many unique barcodes (**Supplementary Table 3**), the few functional barcodes are not expected to impact our measurements, as evidenced by the high correlation observed between cell type replicates in **Fig. 1c** (Methods).

#### Functional 3'UTR element computational modeling

The original MPRAu dataset was first filtered to remove variants that had an average plasmid count of less than 20 (average calculated across all plasmid replicates). For each cell type, for predicting attenuation, oligos were classified as one if their $log_2FC$ was less than -0.5, and zero otherwise. For predicting augmentation, oligos were classified as one if their $log_2FC$ was greater than 0.5, and zero otherwise. We performed predictions independently for each cell type. For prediction, two sets of variables were utilized: the minimal model, and the full model.

For the minimal model, the following variables were calculated across each oligo and used for prediction: nucleotide percentage (across four bases, four variables), dinucleotide percentages (six variables), exact dinucleotide counts (16 variables), minimal free energy (as measured by RNAFold), maximum homopolymer length (for each base type A, U, C, G, and across all base types, five variables), maximum dinucleotide length across all bases, and a measure of sequence uniformity (let "seq" be the tested sequence, then it is calculated as the following: for i in range (1,len (seq),1): if seq[i]==seq[ (i-1)]: seq_uniformity=seq_uniformity+1)

For the full model, all of the variables in the minimal model were used, and the following variables, calculated across the entire oligo, were added per cell type: the number of RBP motifs for AU-rich, GU-rich, CU-rich, GU-rich, constitutive decay, and Pumilio elements (6 variables), RNA Rck scores (Orenstein et al., 2016) for expressed RBPs (79-86 variables total, dependent on which RBPs were expressed in specific cell types), the number of binding sites for each of the top 100 expressed miRNAs within the cell type (100 variables total), the number of hexamers for each of the top 300 most significant activity hexamers within the cell type (ranked by adjusted p-value, 300 variables total), the total number of overlapped miRNA motifs from the top 10/25/100 expressed miRNAs subsetting on 8mer/7mer-1A/7mer-m8/8mer+7mer-1A+7mer-m8/8mer+7mer-m8 annotations (15 variables), and the number of overlapped hexamers derived from the top 10/20/50/100/200/300 most significant positive/negative/all hexamers (ranked by adjusted p-value, 18 variables). We note that all the hexamer and miRNA-associated variables are derived from cell-type-specific datasets, thus are different in each cell type. For the miRNA annotations, if a cell type contained multiple miRNA expression datasets, we used the dataset that had the most significant attenuating effect for oligos that contained motifs from the top 10 most highly expressed miRNAs (subsetting on the 8mer+7mer-m8 annotation, significance was calculated via a *t*-test, with the null mean centered at zero).

For each cell type, under both the minimal and full model tested, the following procedure was performed for training. Approximately 20% of the dataset was used for validating the final model, and ~80% of the dataset was used to train a predictive model using xgboost. The initial ~20% testing was selected by iteratively selecting all the oligos for a specific gene, weighted by the number of oligos corresponding to the gene, until the size of the validation set reached at least 20%. The rest of the oligos that were not selected in the testing set were used in the training set. Some genes had overlapping 3'UTRs, and the oligos that they overlapped were grouped together as being derived from a single gene in this selection strategy. This splitting ensured that oligos in the same gene did not overlap in the validation/training sets.

The xgboost implementation in python was used then used to train models with the following range of parameters (in the following format: [ parameter_name: start, stop, step_size ]): [ colsample_bytree: 0.75, 1, 0.25 ], [ gamma: 1, 7, 2 ], [ min_child_weight: 1, 7, 2 ], [ max_depth: 2, 10, 2 ], and [ n_estimators: 50, 150, 50 ] across all cell types. This range of parameters yielded a total of 480 combinations. We also used SVM and logistic regression, and the implementations in the python sklearn package were used. For parameter tuning, the following parameters were used - for SVM: [ C: 0.0001, 0.001, 0.01, 0.1, 1, 10, 100, 1000 ], [ gamma: 0.001, 0.002, 0.01, 0.02, 0.03, 0.05, 0.1, 1, 2 ], [ kernel: string, rbf ], and for logistic regression: [ C: 0.0001, 0.001, 0.01, 0.1, 1, 10, 100, 1000 ], [ penalty: l1, l2 ]. Lastly, a one-variable decision tree model trained using percent U was used as a control.

For each parameter combination, five-fold cross-validation, grouped by genes (so that oligos will not overlap between the testing and training folds), was performed to obtain the combination's performance. The mean of the average precision across the five folds was utilized as the scoring metric for model performance under each parameter setting. The model parameter combination with the best mean average precision score was chosen as the best combination. To obtain the validation average precision score, the best parameter combination was used to train a model on the entire training set, and then evaluated by the validation set which was never seen in the training procedures. Across all tested cell types, we consistently found xgboost outperforming all models in all predictions (**Supplementary Fig. 4 and 5**).

#### Concordance of allelic skews with Geuvadis and GTEx datasets

Both ASE and eQTL data were downloaded from https://www.ebi.ac.uk/arrayexpress/experiments/E-GEUV-1/files/analysis_results/
Allele-specific expression data was derived from the file GD462.ASE.COV8.ANNOT_PTV.txt.gz and eQTL data was derived from combining the files YRI89.gene.cis.FDR5.all.rs137.txt.gz and EUR373.gene.cis.FDR5.all.rs137.txt.gz.

GM12878 MPRAu variant data was overlapped with Geuvadis ASE data using chromosome, position, ref, and alt allele as the identifier to match between datasets. The overlapped set was then further filtered by only retaining effect sizes which have a p-value less than 0.05 for Geuvadis and a BH p-adj less than 0.1 for MPRAu. Variants were further filtered by keeping the ones that had a pooled (across individuals) Geuvadis ASE significant effect (two-sided *t*-test p <0.01). From the remaining filtered variants, the median Geuvadis ASE value from all individuals per variant was used for plotting in **Fig. 4a**. If multiple MPRAu variants had the same identifier (for example, due to the same variant lying in different backgrounds), the median was taken as the point for plotting.

eQTL data from both YRI and EUR were concatenated together into a single file. GM12878 data was extracted from our MPRAu data set and overlapped with Geuvadis data using chromosome, position, ref, and alt allele, and Ensembl gene name as the identifier to match between datasets. If multiple Geuvadis eQTL entries overlap with a single MPRAu variant, then the median was taken as the representative point for plotting in **Supplementary Fig. 6a**. Similarly, if multiple MPRAu variants had the same identifier (for example, due to the same variant lying in different backgrounds), the median was taken as the point for plotting.

GTEx v8 fine-mapped eQTL effect sizes (Ulirsch) were correlated with MPRAu allelic skews in **Fig. 4b** and **Supplementary Fig. 7a**. Since there was not a direct mapping of MPRA tested cell type to GTEx tissue type, and variants that have significant allelic skew effects across multiple tissues appear to have concordant directional effects across these same tissues (**Fig. 1e**), we created aggregated scores to compare our MPRAu experiments with the GTEx fine-mapping dataset. To do this, for each variant, we took the median eQTL effect size across all GTEx tissues with a fine-mapped signal greater or lesser than a certain PIP cutoff (greater than 0.2 for **Fig. 4b**, and **Supplementary Fig. 7a**, lesser than 0.01 for **Supplementary Fig. 7b**) and compared it against the median tamVar skew across significant skew values (BH p-adj<0.05 for **Fig. 4b** and <0.1 for **Supplementary Fig. 7a**,**b**) from all cell types. We further included only variants that are annotated by the Ensembl Variant Effect Predictor (version 85) (McLaren et al., 2016) to have the 3'UTR annotation as the most severe consequence in comparing allelic effect concordance. MPRAu data was overlapped with the GTEx v8 fine-mapping data using chromosome, position, ref, and alt allele, and Ensembl gene name as the identifier to match between datasets.

#### UK Biobank fine-mapping enrichment

Genetic fine-mapping of 94 traits in up to 361,194 individuals from the UKBB was performed using FINEMAP (Benner et al., 2016) and the Sum of Single Effects (SuSiE) method (Wang et al., 2020a) (https://www.finucanelab.org/data). Fine-mapped variants were overlapped with MPRAu variant data and across all PIP filters, the proportion of tested 3'UTR variants significant in at least one MPRAu tested cell type was used to create **Fig. 4c**. If a variant was tested across multiple sequence backgrounds, it was considered significant if strong allelic effects (BH p-adj<0.1) were observed in any of the tested contexts. We only included variants that were annotated by the Ensembl Variant Effect Predictor (version 85) (McLaren et al., 2016) to have 3'UTR annotation as the most severe consequence. 3'UTR variants that were part of a 95% SuSiE credible set with another highly likely causal variant (defined as PIP>0.3 and in a coding or promoter region) were also excluded.

**3'UTR SNV and deletion tiling analysis**

Log$_2$FC of RNA over plasmid was quantified via DESeq2. To generate the sequence logo for SNV tiling, the sign of the log$_2$FC of the unaltered oligo was first multiplied to the log$_2$FC of all oligos that tiled the specific region. Then, per oligo position, all four log$_2$FC values (corresponding to the position with A, U, C, or G) were normalized by the sum of the total log$_2$FC for that position. These steps created a normalized weight matrix per position across the entire oligo. EDLogo (Dey et al., 2018) then used these matrices to generate the motifs for SNV tiling. In contrast to conventional sequence logos, EDLogo allows both enrichment and depletion of the activity of characters to be displayed at each specific position. Formally, enrichment/depletion for a specific character at a specific position is interpreted to be relative to the median enrichment/depletion across all possible characters for the position.

We multiplied the sign of the log$_2$FC of the unaltered oligo first to all the other oligos to ensure that underlying motifs (regardless of the direction of its effect), if perturbed, show positive enrichment scores. For example, at a specific position, if a base is part of a functional element, and all other bases perturb the element, then the log$_2$FC magnitude observed for the other bases would be less in magnitude than the one observed by the unaltered oligo. In this manner, the base of the unaltered oligo would have the highest enrichment score (as seen in the AU-rich element in **Fig. 5c** and the U1 snRNP motif in **Fig. 5a**). However in the rare case of a single base creating a novel functional element in the opposite effect of the unaltered oligo log$_2$FC, then we would see the base having a very low enrichment score (as seen in the motif derived in the ref SNV tiling plot in **Fig. 5b**).

**Investigating of LD Structure surrounding rs705866 and rs1059273**

The surrounding LD block around rs705866 contains approximately 150 variants, as estimated by looking at the LD block via Haploreg ($r^2$>0.6) (Ward and Kellis, 2016), or the credible set given in Supplementary Data Set 3 from Fritsche et al. 2016. Most noticeably, amongst these ~150 variants, three are missense variants. Although the three missense SNPs are in LD (rs11771799, rs35986051, and rs11761306) with rs705866, none of them look to be strongly conserved (**Supplementary Fig. 9a**), and all of them are also annotated as benign according to the Ensembl Variant Effect Predictor (McLaren et al., 2016). Examining the protein structure of *PILRB*, all three also lie on non-secondary structure elements, far away from the active site for the gene, which further gives support of these variants being non-functional (**Supplementary Fig. 9b**). While another variant may still have causal regulatory effects, it is still likely that rs705866 is a causal variant due to the concordance in MPRAu/HDR allelic expression directionality with the reported retina eQTL allelic effect (i.e. higher expression in the alt allele).

For rs1059273, FsT was calculated between all pairwise non-admixed populations in the 1000 Genomes Dataset (1000 Genomes Project Consortium et al., 2015) using the StAMPP package (https://www.rdocumentation.org/packages/StAMPP/versions/1.5.1/topics/StAMPP-package) (Pembleton et al., 2013). The highest significantly elevated fixation index was observed between Han Chinese and Esan. These populations were then used to calculate EHH scores on ancestral and derived haplotypes centered at rs1059273. Related individuals were excluded from all analyses.

### RESOURCE AVAILABILITY

#### Lead Contact
Further information and requests for resources and reagents should be directed to James Xue.

#### Materials Availability
Oligo libraries used in this study are available upon request. CRISPR-modified GM12878s for rs1059273 and CRISPR-modified SK-N-SH for rs705866 are available upon request. All additional unique/stable reagents generated in this study are available from the Lead Contact without restriction, or with a Materials Transfer Agreement.

#### Data and Code Availability
Raw sequencing reads are being deposited to GEO, the ENCODE portal (https://www.encodeproject.org/), and SRA. Processed MPRAu screen data are available in **Supplementary Table 1** and will also be available on the ENCODE portal. Read counts per oligo are included in **Supplementary Table 1**. Analyses was performed with standard analysis packages cited in the text, and plotted using custom R scripts that are available upon request.

#### Key Resource Table

| REAGENT or RESOURCE | SOURCE | IDENTIFIER |
|---|---|---|
| Bacterial and Virus Strains | | |
| 10-beta Electrocompetent E. coli | NEB | C3020K |
| Chemicals, Peptides, and Recombinant Proteins | | |
| Q5 NEBNext MasterMix | NEB | M0541S |
| Q5 HotStart polymerase | NEB | M0493S |
| BSA | NEB | B9000 |
| NEBNext Ultra II Q5 Master Mix | NEB | M0544L |
| Agencourt AMPure XP SPRI | Beckman Coulter | A63881 |
| Gibson Assembly Master Mix | NEB | E2611 |
| BmtI | NEB | R0658S |
| XbaI | NEB | R0145S |
| SOC | NEB | B9020S |
| Kanamycin | Teknova | K2125 |
| DMEM | ThermoFisher | 10564 |
| FBS | ThermoFisher | A3160501 |
| MEM Non-Essential Amino Acids Solution | ThermoFisher | 11140050 |
| Sodium Pyruvate | ThermoFisher | 11360070 |
| Lipofectamine 2000 Transfection Reagent | ThermoFisher | 11668027 |
| Cycloheximide Solution | Sigma-Aldrich | C4859 |
| Tris-HCl | ThermoFisher | 15567 |
| $MgCl_2$ | ThermoFisher | AM9530G |
| KCl | ThermoFisher | AM9640G |
| Dithiothreitol | VWR | 97061-340 |
| Triton X-100 | Sigma-Aldrich | T8787 |
| Sodium Deoxycholate | Sigma-Aldrich | 30970 |
| NaCl | ThermoFisher | AM9760G |
| RPMI 1640 Medium, GlutaMAX Supplement | ThermoFisher | 61870036 |
| Penicillin-Streptomycin | ThermoFisher | 15140122 |
| Corning Penicillin-Streptomycin Solution | Corning | 30-002-CI |

| | | |
|---|---|---|
| MEM α, GlutaMAX Supplement | ThermoFisher | 32561037 |
| Lipofectamine 3000 Transfection Reagent | ThermoFisher | L3000015 |
| EMEM | ATCC | 30-2003 |
| MEGM BulletKit | Lonza | CC-3150 |
| TRIzol LS Reagent | ThermoFisher | 10296 |
| GlycoBlue Coprecipitant | ThermoFisher | AM9516 |
| Oligo d(T)$_{25}$ Magnetic Beads | NEB | S1419S |
| TURBO DNase | ThermoFisher | AM2239 |
| SDS | ThermoFisher | AM9822 |
| Agencourt RNAClean XP SPRI | Beckman Coulter | A63987 |
| SuperScript III Reverse Transcriptase | ThermoFisher | 18080 |
| SYBR Green I Nucleic Acid Gel Stain | Life Technologies | S-7567 |
| High Sensitivity D1000 Reagents | Agilent | 5067-5585 |
| Power SYBR Green PCR Master Mix | ThermoFisher | 4367659 |
| Alt-R CRISPR-Cas9 tracrRNA | IDT | 1072533 |
| Nuclease-Free Duplex Buffer | IDT | 1072570 |
| Alt-R S.p. Cas9 Nuclease V3 | IDT | 1081059 |
| Alt-R Cas9 Electroporation Enhancer | IDT | 1075916 |
| Alt-R HDR Enhancer | IDT | 1081072 |
| Alt-R A.s. Cas12a (Cpf1) Ultra | IDT | 10001273 |
| Alt-R Cpf1 Electroporation Enhancer | IDT | 1076300 |
| Critical Commercial Assays | | |
| Dual-Glo Luciferase Assay System | Promega | E2920 |
| QIAGEN Plasmid Plus Giga Kit | QIAGEN | 12991 |
| Neon Transfection System 100 µL Kit | ThermoFisher | MPK10096 |
| Neon Transfection System 10 µL Kit | ThermoFisher | MPK1025 |
| AllPrep DNA/RNA Mini Kit | QIAGEN | 80204 |
| Deposited Data | | |
| Raw Sequencing Reads (in process of deposit) | GEO, SRA | NA |
| Processed MPRAu Data | ENCODE | NA |
| Experimental Models: Cell Lines | | |
| HEK293FT | ThermoFisher | R70007 |
| HepG2 | ATCC | HB-8065, RRID:CVCL_0027 |
| HMEC | ThermoFisher | A10565 |
| GM12878 | Coriell | GM12878, RRID:CVCL_7526 |
| K562 | ATCC | CCL-243, RRID:CVCL_0004 |
| SK-N-SH | ATCC | HTB-11, RRID:CVCL_0531 |
| CRISPR-modified GM12878s for rs1059273 | NA | NA |
| CRISPR-modified SK-N-SH for rs705866 | NA | NA |
| Oligonucleotides | | |
| Primers | IDT | NA |
| ssODN | IDT | NA |
| CMS array oligos | CustomArray | NA |
| GWAS array oligos | Twist Biosciences | NA |
| Alt-R CRISPR-Cas9 tracrRNA | IDT | 1072533 |
| Recombinant DNA | | |
| pmirGLO vector | Promega | E1330 |

| | | |
|---|---|---|
| pGL4.23[luc2/minP] Vector | Promega | E8411 |
| Software and Algorithms | | |
| DESeq2 | https://bioconductor.org/packages/release/bioc/html/DESeq2.html | RRID:SCR_015687 version 1.22.2 |
| ggplot2 | https://cran.r-project.org/web/packages/ggplot2/index.html | RRID:SCR_014601 version 3.1.1 |
| Other | | |
| Nunc TripleFlask | VWR | 89498-706 |

**Supplementary Tables:**
**Supplementary Table 1: MPRAu results**

**Supplementary Table 2: Luciferase assay results**

**Supplementary Table 3: tamVars with gene expression and phenotype fine-mapping support**

**Supplementary Table 4: MPRAu barcode effects**

**Supplementary Table 5: Primers, HDR ssODN/guides**

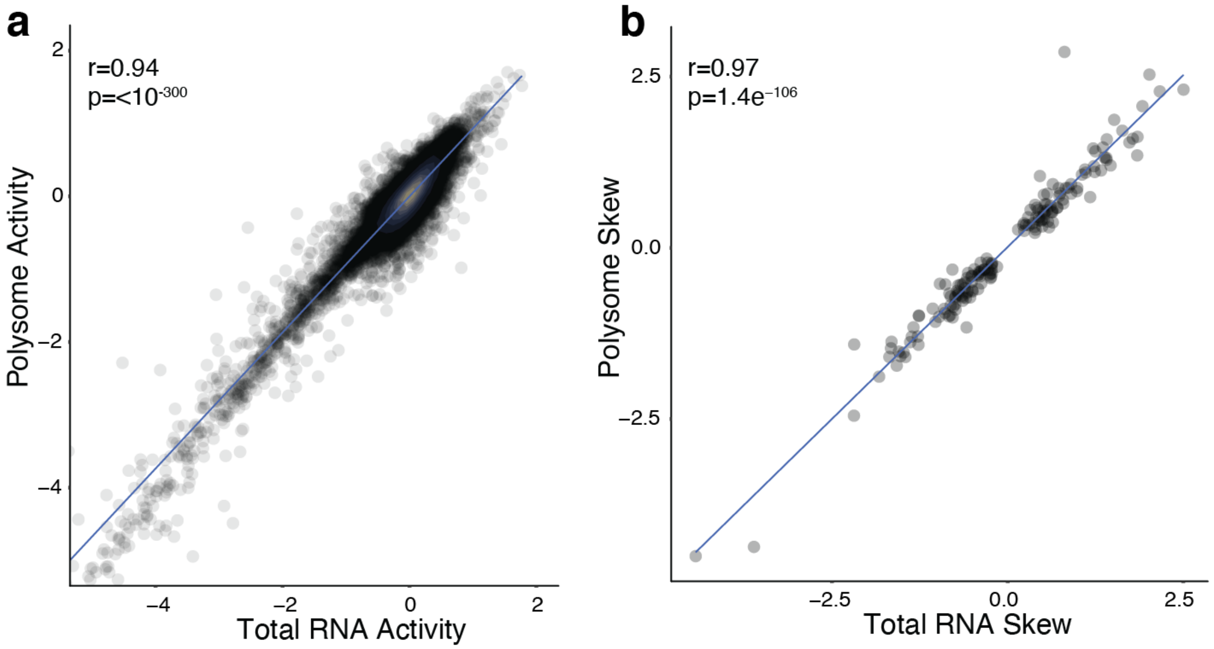


**Supplementary Fig. 1: Polysomal RNA-derived MPRAu metrics are correlated with Total RNA-derived MPRAu metrics.**

**a**, HEK293 Polysomal RNA-derived MPRAu activity is correlated with total RNA-derived HEK293 MPRAu activity. **b**, HEK293 Polysomal RNA-derived MPRAu skew is correlated with HEK293 total RNA-derived MPRAu skew for significant hits (BH p-adj<0.1 for either Polysomal RNA or total RNA variants).

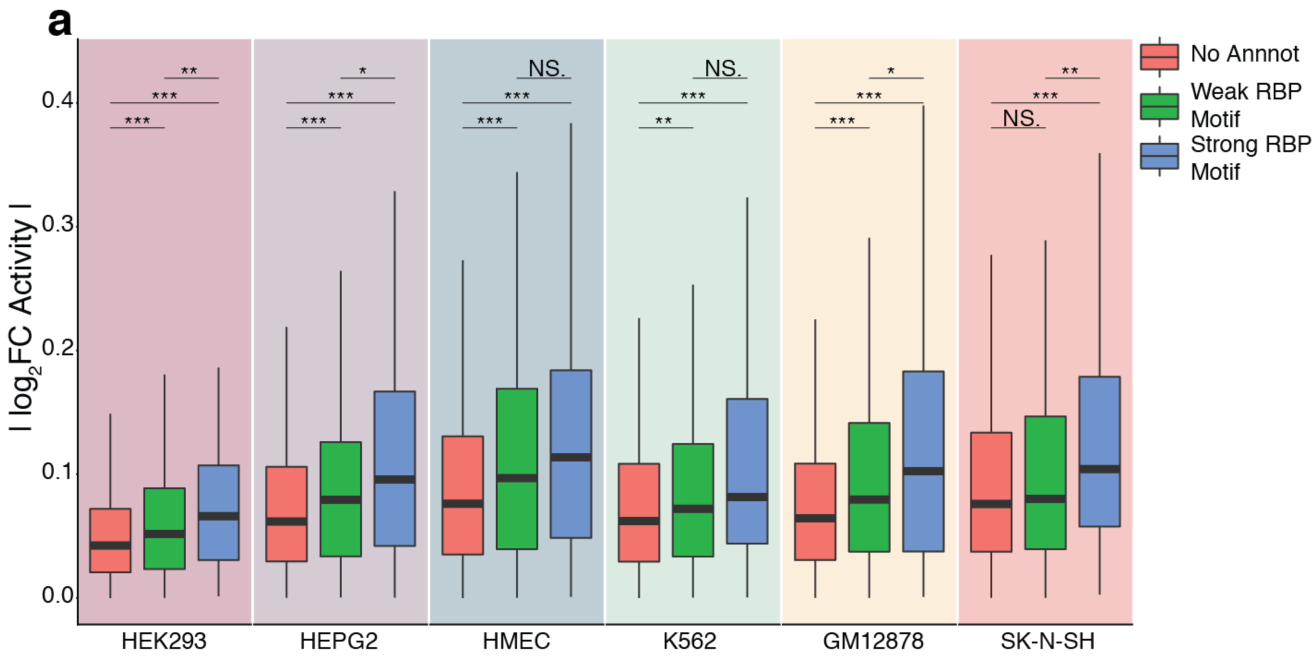


**Supplementary Fig. 2: MPRAu hexamer barcodes demonstrate activity across hexamers with known functionality.**

**a**, MPRAu barcodes that match RBP motifs derived from Dominguez et al., 2018 have a greater magnitude in 3'UTR activity.

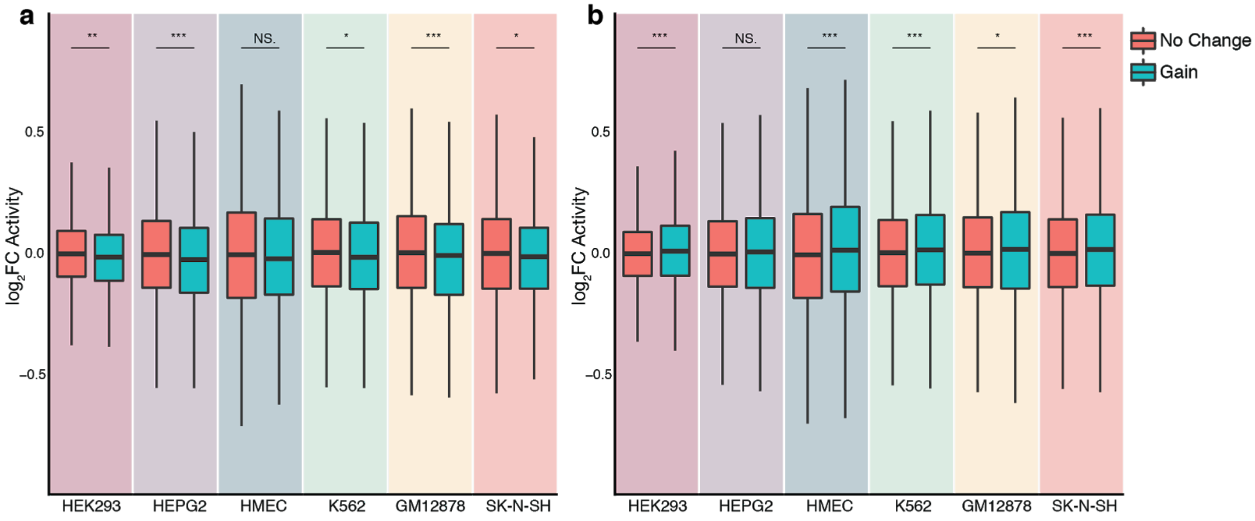


**Supplementary Fig. 3: Gain of hexamer barcodes with directional activities leads to increased MPRAu-measured effects.**

**a**, Gain of a top 20 barcode with negative activity (ranked by p-adj) leads to attenuation across all tested cell types. **b**, Gain of a top 200 barcode with positive activity (ranked by p-adj) leads to augmentation across all tested cell types.

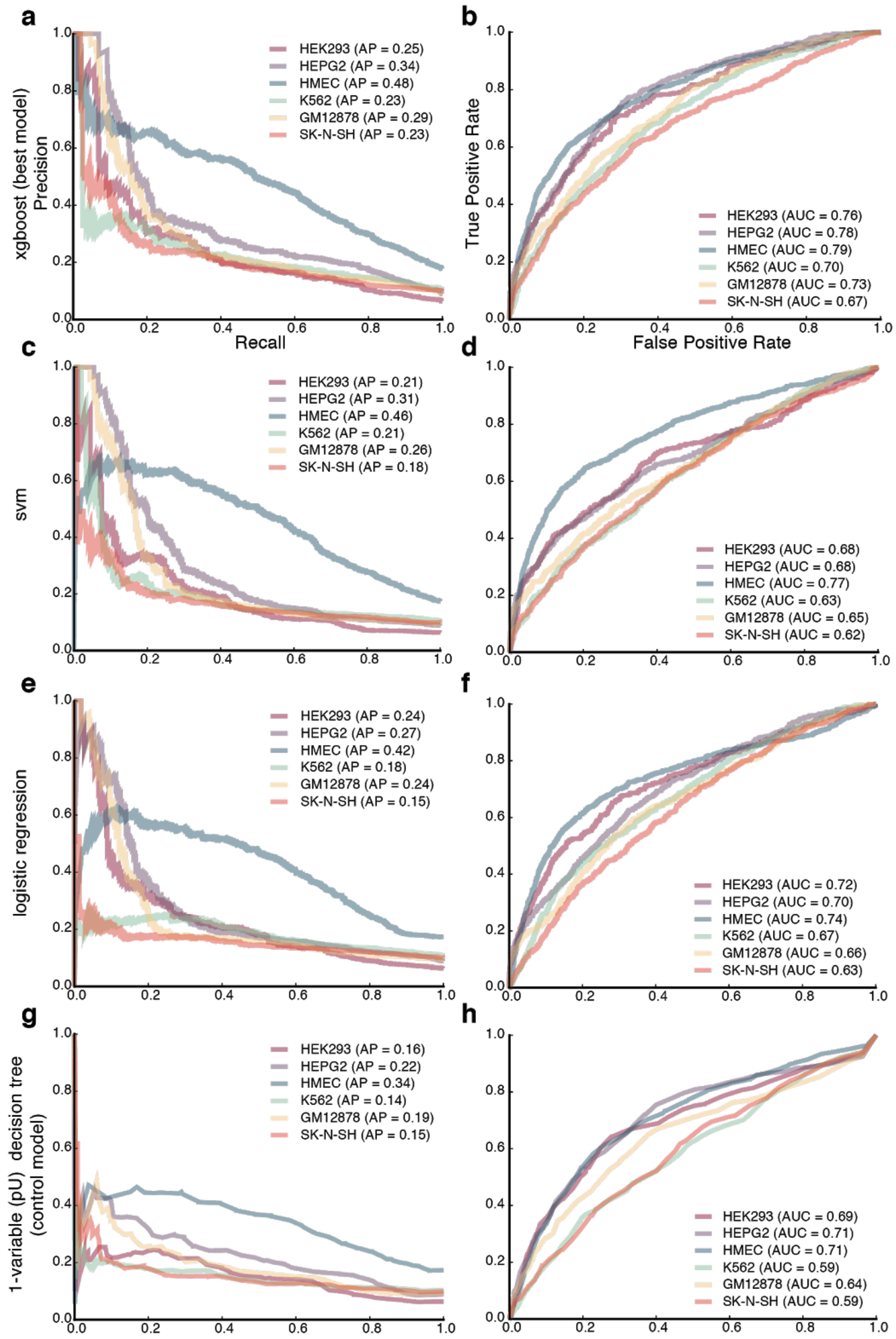


**Supplementary Fig. 4: xgboost outperforms other prediction methods across all cell types in predicting attenuation using sequence-specific annotations.**

**a**, xgboost precision-recall curve across all six tested cell types. **b**, xgboost ROC curve across all six tested cell types. **c**, SVM precision-recall curve across all six tested cell types. **d**, SVM ROC curve across all six tested cell types. **e**, Logistic regression precision-recall curve across all six tested cell types. **f**, Logistic regression ROC curve across all six tested cell types. **g**, One-variable percent U decision tree precision-recall curve across all six tested cell types. **h**, One-variable percent U decision tree ROC curve across all six tested cell types.

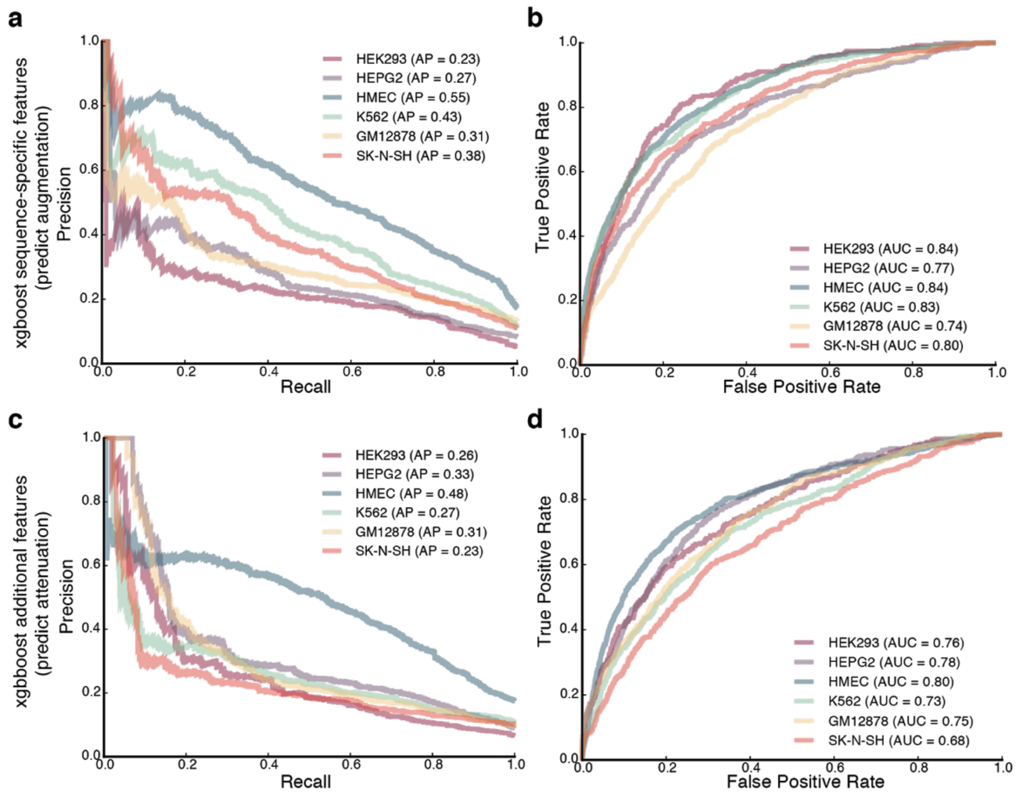


**Supplementary Fig. 5: Inclusion of additional variables for xgboost does not improve attenuation prediction, and xgboost can predict MPRAu oligos with augmenting activity as well.**

**a**, xgboost precision-recall curve across all six tested cell types for predicting augmenting activity. **b**, xgboost ROC curve across all six tested cell types for predicting augmenting activity.

**c**, xgboost precision-recall curve across all six tested cell types for the full model in predicting attenuation.

**d**, xgboost ROC curve across all six tested cell types for the full model in predicting attenuation.

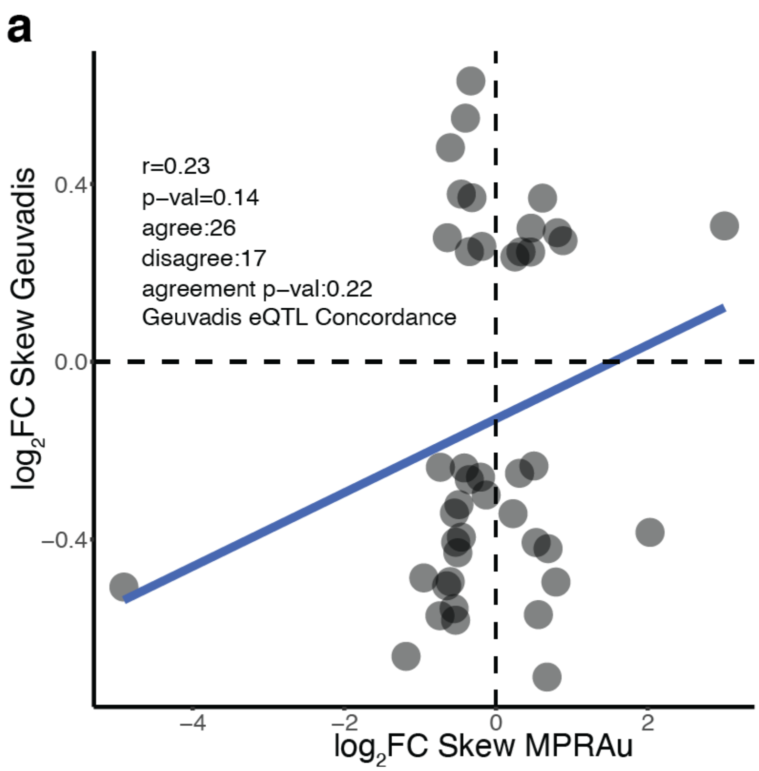


**Supplementary Fig. 6: Geuvadis eQTLs are less concordant with 3'UTR activity.**
**a**, Geuvadis eQTL has a positive, yet non-significant trend, with GM12878 3'UTR activity scores.

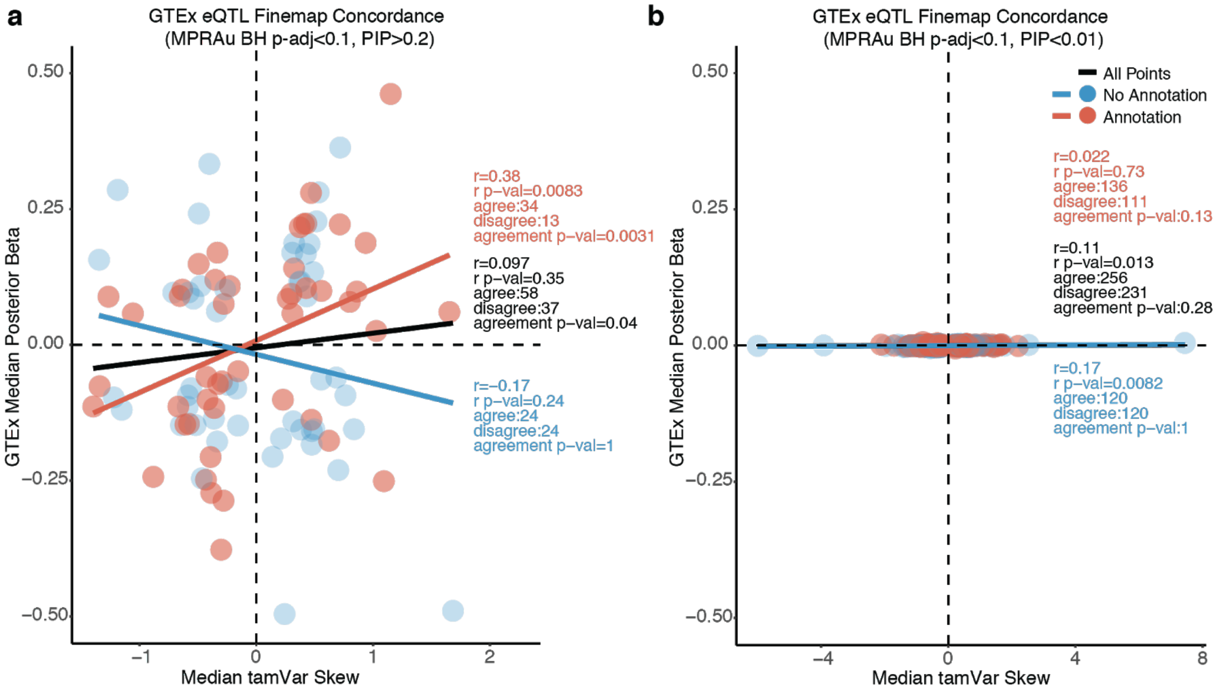


**Supplementary Fig. 7: GTEx fine-mapping concordance with MPRAu results**
**a**, Relaxing MPRAu BH p-adj cutoff to 0.1 still yields concordance between GTEx and MPRAu effects. **b**, Concordance is lost at low a PIP cutoff (<0.01).

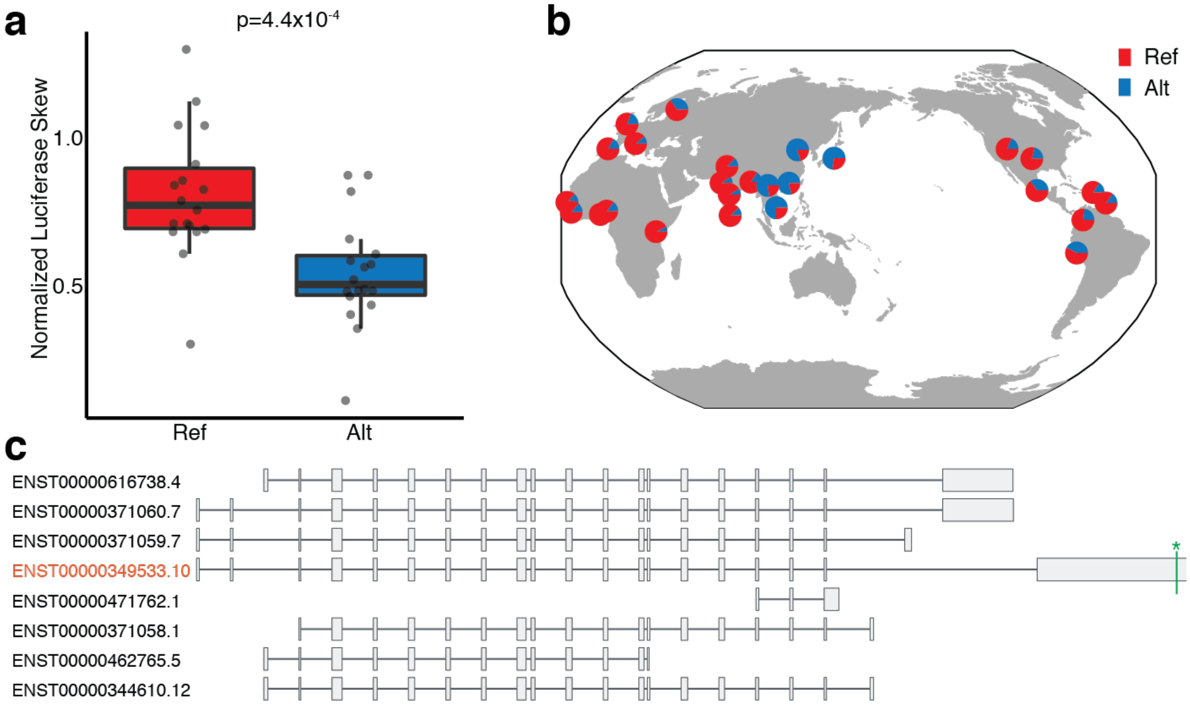


**Supplementary Fig. 8: Additional information relevant to rs34448361**
**a**, Luciferase assay of rs34448361 validates the tamVAR effect observed in MPRAu. **b**, The alternate allele (insertion that extends the AU-rich element overlying the variant) has high frequency specifically in East Asian populations. **c**, rs34448361(green star) lies specifically in the 3'UTR of the longest isoform of *LEPR* ENST00000349533 (*LepRb*).

**a**

| | | | | | | | | rs11771799 (T to C) | | | rs35986051 (T to C) | | | | | rs11761306 (A to G) | | | | |
|---|---|---|---|---|---|---|---|---|---|---|---|---|---|---|---|---|---|---|---|---|
| Human | T | T | A | G | C | C | A | T | A | G | T | T | C | C | C | A | A | C | G | T |
| Chimp | T | T | A | G | C | C | A | C | A | G | C | T | C | C | C | G | A | C | G | T |
| Bonobo | T | T | A | G | C | C | A | C | A | G | C | T | C | C | C | G | A | C | G | T |
| Gorilla | T | T | A | A | C | C | A | C | A | G | C | T | C | C | C | G | A | T | G | T |
| Orangutan | T | T | A | G | C | C | A | C | A | G | C | T | T | C | C | A | A | C | A | T |
| Gibbon | T | T | A | G | C | C | A | C | A | G | C | T | C | C | T | A | A | C | A | T |
| Rhesus | T | T | A | G | C | C | G | C | A | G | C | T | C | C | C | A | A | C | A | T |
| Crab-eating macaque | T | T | A | G | C | C | G | C | A | G | C | T | C | C | C | A | A | C | A | T |
| Pig-tailed macaque | T | T | A | G | C | C | G | C | A | G | C | T | C | C | C | A | A | C | A | T |
| Sooty mangabey | T | T | A | G | C | C | A | C | A | G | C | T | C | C | C | A | A | C | A | T |
| Baboon | T | T | A | G | C | C | A | C | A | G | C | T | C | C | C | A | A | C | A | T |
| Green monkey | T | T | A | G | C | C | A | C | A | G | C | T | C | C | C | A | A | C | A | T |
| Drill | T | T | A | G | C | C | A | C | A | G | C | T | C | C | C | A | A | C | A | T |
| Angolan colobus | T | T | A | G | C | C | A | C | A | G | C | T | C | C | C | A | A | C | A | T |
| Golden snub-nosed monkey | T | T | A | G | C | C | A | C | A | G | C | T | C | C | C | A | A | C | A | T |
| Black snub-nosed monkey | T | T | A | G | C | C | A | C | A | G | C | T | C | C | C | A | A | C | A | T |
| Marmoset | T | T | A | G | C | C | A | G | A | C | C | T | C | C | G | A | A | C | A | T |
| Squirrel monkey | T | T | A | G | C | C | A | G | A | C | C | T | C | C | C | A | A | T | G | T |
| White-faced sapajou | T | T | A | G | C | C | A | G | A | C | C | T | C | C | C | A | A | T | G | T |
| Ma's night monkey | T | T | A | G | C | C | A | G | A | C | C | T | C | C | C | A | A | T | G | T |
| Tarsier | T | T | A | G | C | C | G | A | G | G | A | T | C | C | C | A | A | A | G | T |
| Mouse lemur | T | T | A | G | C | C | A | A | A | G | A | T | C | C | C | A | A | C | A | T |
| Coquerel's sifaka | T | T | A | G | C | C | A | A | A | G | T | T | C | C | C | A | G | A | G | T |
| Black lemur | T | T | A | G | C | C | A | A | A | G | T | T | C | C | C | A | A | T | G | T |
| Sclater's lemur | T | T | A | G | C | C | A | A | A | G | T | T | C | C | C | A | A | T | G | T |
| Bushbaby | T | T | A | G | C | T | G | A | - | - | - | T | C | C | C | A | G | A | G | T |
| Dog | T | T | A | G | C | C | A | A | G | G | A | T | C | C | C | A | G | A | G | T |
| Armadillo | T | T | G | G | C | C | A | A | G | G | A | G | C | C | C | A | G | G | C | T |
| Proboscis monkey | = | = | = | = | = | = | = | = | = | = | = | = | = | = | = | = | = | = | = | = |

**b**

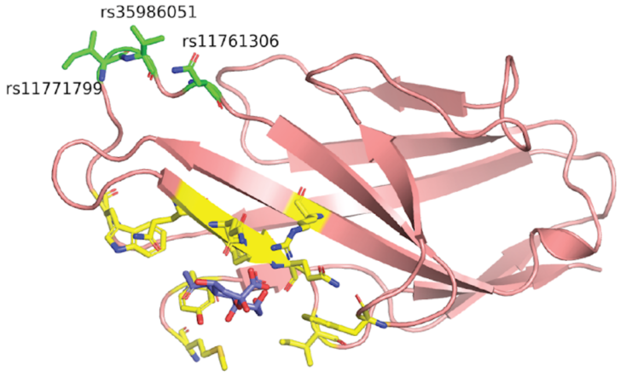


**Supplementary Fig. 9: Missense SNPs in LD with rs705866 are predicted to be benign**
**a**, Multiple sequence alignment across 30 mammals show that rs11771799, rs35986051, and rs11761306 (all in LD with rs705866) are in bases without strong conservation. **b**, Protein structure of PILRB demonstrates that rs11771799, rs35986051, and rs11761306 (green) lie far from the active site and are likely non-functional mutations. Shown in yellow are residues within 5 Å of the sialic acid ligand (purple).

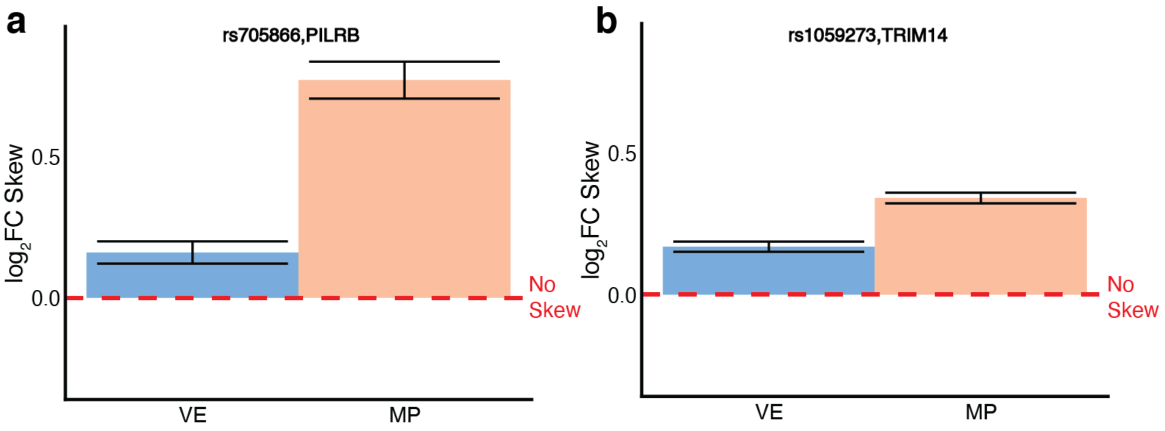


**Supplementary Fig. 10: Replication of allelic replacement effects for *PILRB* and *TRIM14.***
**a**, Allelic skew effects from the second replicate of a HDR allelic replacement of rs705866 in SK-N-SH recapitulates the result seen in the first replicate (VE: Fisher $p=1.43 \times 10^{-15}$, MP: Fisher $p=2.53 \times 10^{-116}$). **b**, Allelic skew effects from the second replicate of a HDR allelic replacement of rs1059273 in GM12878 recapitulates the result seen in the first replicate (VE: Fisher $p=5.40 \times 10^{-76}$, MP: Fisher $p=1.83 \times 10^{-287}$).
